# A novel mechanobiological model of bone metastasis reveals that mechanical stimulation inhibits the pro-osteoclastogenic effects of breast cancer cells

**DOI:** 10.1101/2022.09.19.508588

**Authors:** Vatsal Kumar, Syeda M. Naqvi, Anneke Verbruggen, Eoin McEvoy, Laoise M. McNamara

**Author notes:** Both authors contributed to this manuscript equally (Vatsal Kumar, Syeda M. Naqvi).

## Abstract

Bone is highly susceptible to cancer metastasis, and both tumour and bone cells enable tumour invasion through a “vicious cycle” of biochemical signalling. Tumor metastasis into bone also alters biophysical cues to both tumour and bone cells, which are highly sensitive to their mechanical environment. However, the mechanobiological feedback between these cells that perpetuates this cycle has not been studied. Here, we develop highly novel in vitro and computational models to provide an advanced understanding of how tumor growth is regulated by the synergistic influence of tumour-bone cell signaling and mechanobiological cues. In particular, we develop the first multicellular healthy and metastatic bone models, which can account for physiological mechanical signals within a custom bioreactor. These models successfully recapitulated mineralization, mechanobiological responses, osteolysis and metastatic activity. Ultimately we demonstrate that mechanical stimulus provided protective effects against tumor-induced osteolysis, confirming the importance of mechanobiological factors in bone metastasis development.

## 1. Introduction

Tumor and bone cells interact to enable cancer metastasis into bone as part of a complex process known as the ‘vicious cycle’^1, 2^. Osteoblasts release chemokines that facilitate homing of tumor cells to bone^3^, whereas tumor cells produce biochemicals and growth factors (PTHrP, matrix metallopeptidases, interleukins, VEGF) that stimulate osteoclasts to degrade bone matrix^4^. This resorption releases growth factors (TGF-β, IGFs) that stimulate further tumor cell proliferation, thereby perpetuating tumor and bone cell activity^4^. Although the pathogenesis of metastatic bone disease has been widely studied, it remains that therapies are deficient and once cancer invades bone tissue the condition is widely untreatable. Thus, there is a distinct need to significantly advance our scientific understanding of bone metastasis.

Mechanobiological factors have been implicated in breast cancer invasion, homing, tumor growth and metastasis^5–12^. Within the primary tumor, changes in the extracellular matrix composition occur and alter the biophysical environment^13^, which can promote proliferation and transition of tumor cells to migrating cells^8–10, 14–18^. Breast cancer cells become more proliferative, express metastatic genes, and promote osteolysis when cultured on high stiffness substrates ^19, 20^. As they evolve, metastatic bone tumors become less stiff compared to non-tumor bone tissue^21, 22^ and can exert tumor pressure, which likely alters the biophysical cues acting on bone cells. Given the well-established relationship between matrix properties, mechanical stimuli and bone biology, such changes may activate mechanobiological responses in both native bone cells, and also tumor cells, perpetuating the “vicious cycle” driving bone lysis and tumor growth. However, how the coupled mechanobiological responses of tumor and bone cells contribute to osteolysis and tumor growth is not understood.

Much of our understanding of bone metastasis comes from animal models^23–28^, in which normal immune responses are suppressed, or in vitro 2D cell culture studies, which cannot represent the complexity of in vivo cell morphology and interactions with their surrounding matrix^29^. Three-dimensional (3D) in vitro tumor models have been developed to study the invasion of cancer cells into human bone samples. These approaches have included ex-vivo culture of breast cancer cells on human bone tissue ^30, 31^, bio-printed scaffolds cellularized with human bone cells and prostate or breast cancer cells^32, 33^, or encapsulation of bone, endothelial and breast cancer cells in a fibrin hydrogel^34^. Such models have demonstrated the ability of breast cancer cells to colonize various biomaterial and bone scaffolds and the application of such models to study organ-tropism of cancer cells towards bone and sensitivity to chemotherapy drugs. Mineralized collagen matrices and decellularized bone scaffolds have demonstrated the importance of matrix mineralization for inhibiting breast cancer cell proliferation, which led to quiescence through a reduction in mechanosignalling^35^. More recently, a 3D in vitro tumor model was developed to study the invasion of mammary carcinoma cells encapsulated in hydrogels of different substrate stiffness and subjected to a cyclic mechanical compression in a custom bioreactor ^36^. This study revealed that tumor spheroid growth was restricted in stiffer and mechanically compressed hydrogels^36^. Indeed, mechanical compression inhibited tumor growth and osteolysis both in vivo^37, 38^ and in vitro^38^. However, the role of mechanical stimulation in the bone environment on the development of osteolytic bone metastases is not yet fully understood. To address this question requires advanced experimental models that can replicate both the complex multicellular niche and the native mechanical environment that exist in vivo^37^.

In this study, we aimed to develop an advanced in vitro model that can provide an effective surrogate of the metastatic in vivo environment. We first investigated the stress-dependence of tumor spheroid growth through the development of 3D in vitro and computational models. We then extended these models to study the coupled influence of bone cell signaling and mechanical loading on tumor growth. Next, we developed a novel multicellular (osteoblasts, osteocytes, osteoclasts) and mineralised bone model, incorporating mechanical loading using a custom designed bioreactor. Finally, we further developed this model to study metastasis by breast cancer cells and applied this novel model to investigate the influence of mechanical loading on osteolytic activity.

## 2. Results

### 2.1 Mechanical loading reduces tumor spheroid size by restricting cell proliferation

To investigate the mechanoregulation of spheroid expansion, we first encapsulated 4T1 breast cancer cells (in monoculture) within gelatin-transglutaminase hydrogels of varying compression modulus (Figure 1a). We monitored spheroid development over 7 days. Starting from single cells, tumor cells grew into spheroids during the culture period, whose size decreased with increasing hydrogel stiffness (Figure 1c, d). Such changes in size were associated with a reduction in the number of cells in stiffer hydrogels (Figure 1b).

**Figure 1:**
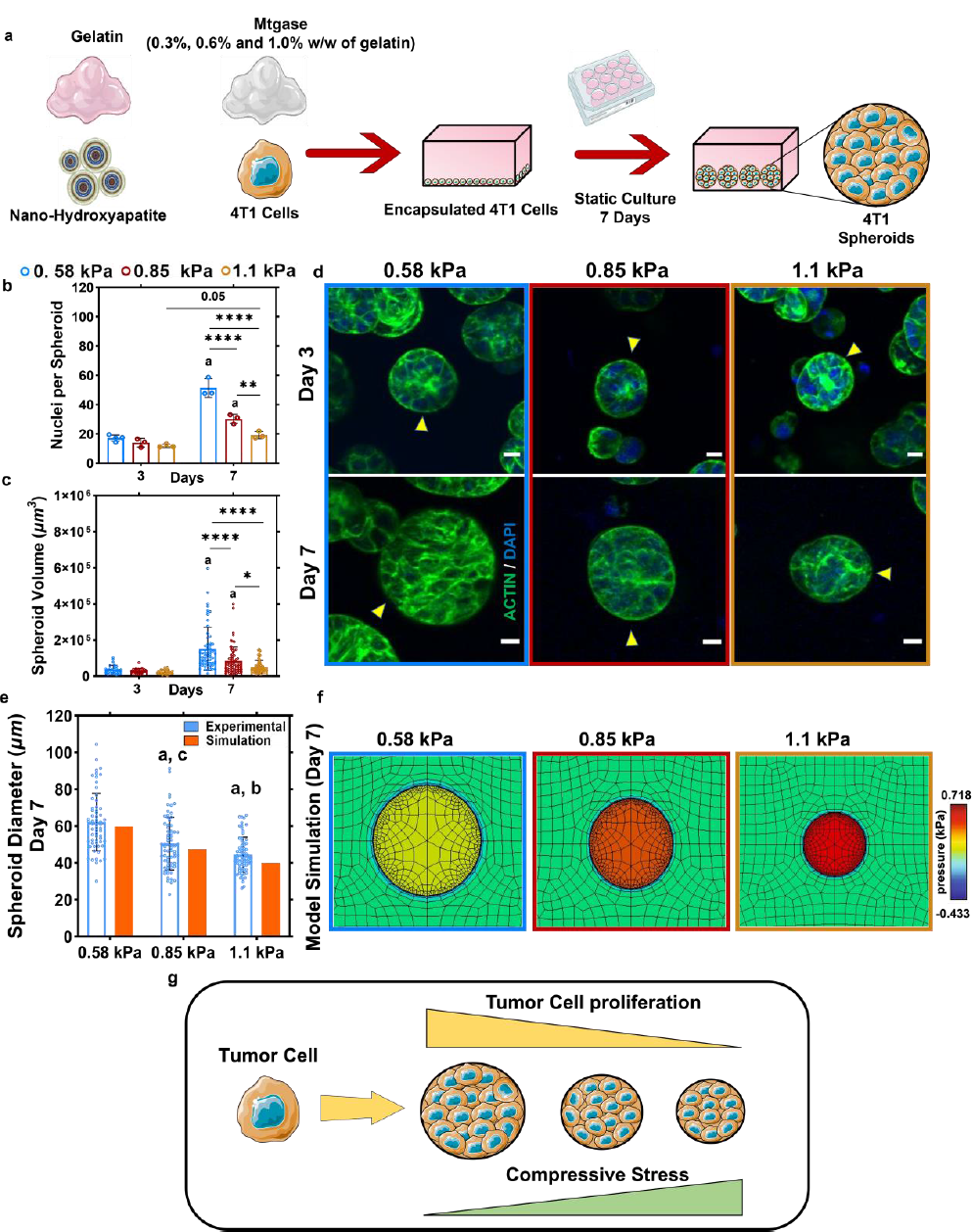
Mechanical loading reduces tumor spheroid size by restricting cell proliferation. (**a**) Schematic of the process of encapsulation of 4T1 cells in single suspension within gelatin-transglutaminase hydrogels and static culture and growth of tumor cells into spheroids. (**b**) Nuclei per tumor spheroid and (**c**) spheroid volume over time across three compression moduli **(0.58 kPa, 0.85 kPa and 1.1 kPa)**. (**d**) fluorescently stained 4T1 spheroids (arrow) within the hydrogels for Actin (green) and DAPI (blue) at day 7 (scale bar: 10 μm). (**e**) Comparison of tumor spheroids in the hydrogels at day 7 of culture with results of simulation from computational tumor growth model within the hydrogels of 0.58 kPa, 0.85 kPa and 1.1 kPa compression moduli. (**f**) Computational model simulation of spheroid growth within the hydrogels (represented by the green background) the resulting stresses that develop. (**g**) schematic describing the relationship between the growth of a single tumor cell into spheroid and the stiffness of the substrate it is confined in. #, a, b, c: significance compared to 4T1 monoculture, 0.58 kPa, 0.85 kPa and 1.1 kPa, respectively.

We next developed a computational model to characterize stress-dependent growth and investigate whether growth was driven by cellular mechanosensitivity (see Methods Section 5.13). The results from this model revealed that increased circumferential and radial stresses, generated by spheroid expansion in a stiffer microenvironment, can reduce the rate of cell proliferation (see Supplementary Figure 1), in agreement with our experimental observations (Figure 1e, f). Altogether, while there may be compaction due to the increased stiffness, we present clear evidence of mechanosensitive reduction in proliferation of the tumor cells in a stiff environment.

### 2.2 Osteoclast precursors have an inhibitory effect on tumor spheroid growth, which is partially alleviated in the presence of osteoblasts

We next explored the coupled influence of hydrogel stiffness and bone cell signaling on in vitro tumor development. For this purpose, we encapsulated osteoclast precursor cells (RAW264.7) and osteoblast-like cells (MC3T3-E1) within separate gelatin-transglutaminase hydrogel suspensions and cultured them in different configurations (monoculture (4T1), co-culture (RAW264.7+4T1, MC3T3+4T1) or tri-culture (MC3T3+RAW264.7+4T1)) with 4T1 cells for 7 days (Figure 2a).

**Figure 2:**
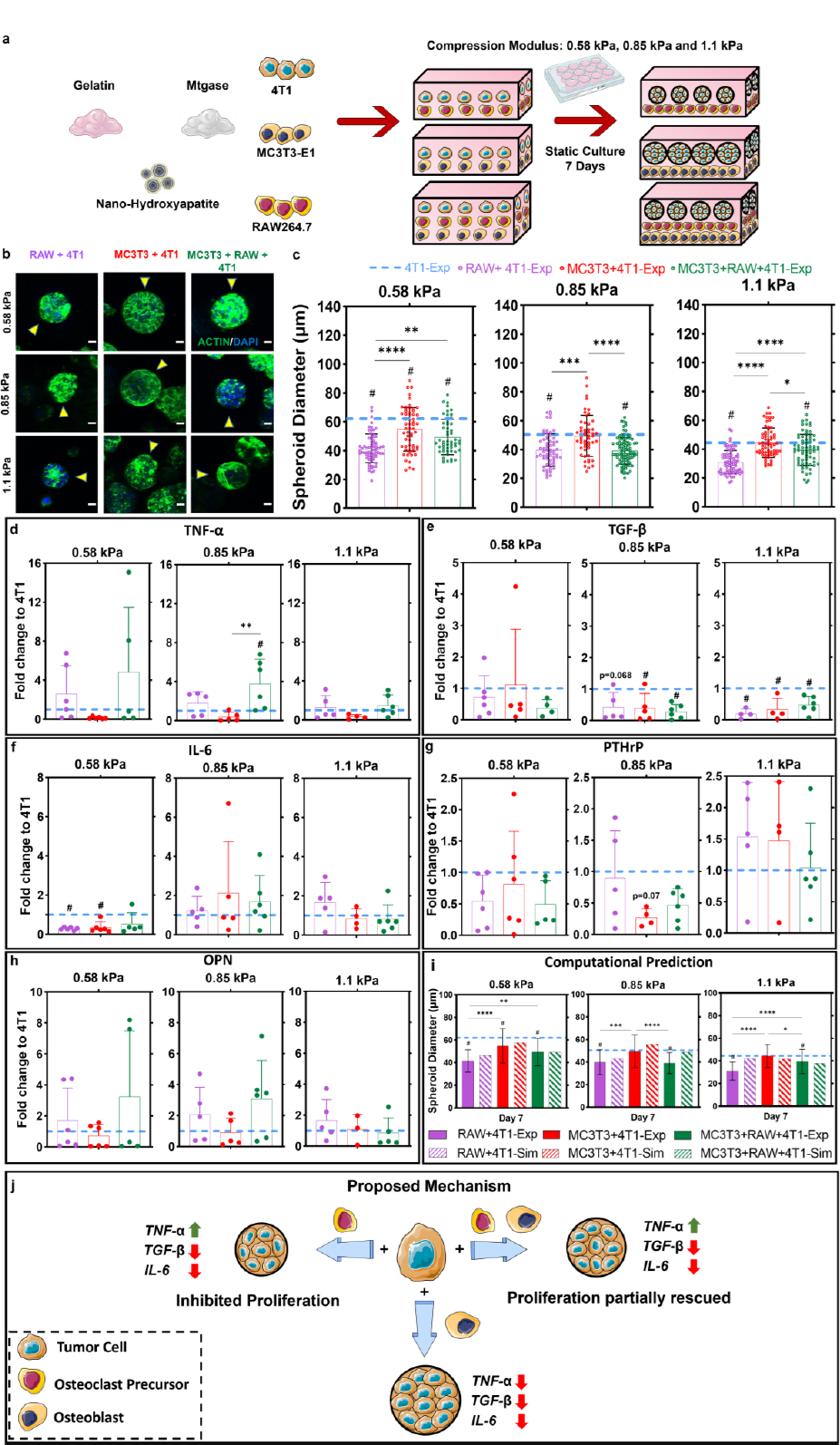
Osteoclasts have an inhibitory effect on 4T1 spheroid growth, which is partially mitigated by osteoblasts. (**a**) Process of encapsulation of RAW264.7 osteoclast precursors, MC3T3-E1 osteoblasts and 4T1 cells within gelatin-transglutaminase hydrogels followed by static co-culture or tri-culture and spheroid formation. (**b**) Fluorescent staining of actin cytoskeleton (green) and nucleus (blue) in 4T1 spheroids (yellow arrows) at day 7 (scale bar: 10 μm). (**c**) Quantification of 4T1 spheroid sizes. (**d-h**) qRT-PCR analysis for TNF-α, TGF-β, IL-6, PTHrP and OPN. (**i**) Computational predictions (Sim) compared to experimental (Exp) tumor spheroid diameter (day 7). (**j**) Proposed biochemical interaction between bone and tumor cells influencing tumor spheroid growth. # significance compared to 4T1 monoculture (blue dashed line) *: p ≤ 0.05, **: p ≤ 0.01, ***: p ≤ 0.001, ****: p ≤ 0.0001.

In the MC3T3 co-culture (MC3T3+4T1), at day 7 of culture, the size of the 4T1 tumor spheroids was significantly smaller relative to those formed by 4T1 cells in monoculture, but only in the 0.58 kPa hydrogels (Figure 2b, c). We observed no difference in the spheroid sizes between the 4T1 monoculture and MC3T3 co-culture groups for the 0.85 kPa and 1.1 kPa hydrogels. However, when co-cultured with RAW264.7 cells (RAW264.7+4T1), the tumor spheroids were significantly smaller than those formed in monoculture for the 0.58 kPa, 0.85 kPa and 1.1 kPa hydrogels (Figure 2b, c). This effect was partially mitigated by the inclusion of MC3T3 cells (i.e., tri-culture group) in 0.58 and 1.1 kPa hydrogels (Figure 2b, c).

Next, we sought to understand tumor spheroid growth in the presence of osteoblasts and osteoclast precursors by investigating expression of genes implicated in tumor growth (TNF-α, TGF-β, OPN) and biochemical signaling between bone and tumor cells (IL-6 and PTHrP). ***Tumor growth:*** The expression of TGF-β was significantly downregulated in the MC3T3 co-culture, and tri-culture groups, compared to the 4T1 monoculture for the 0.85 kPa hydrogels. This effect was also observed for RAW264.7 co-culture, MC3T3 co-culture and the tri-culture groups in the 1.1 kPa hydrogels (Figure 2e). Interestingly, TNF-α expression was significantly upregulated in the tri-culture group when compared to 4T1 monoculture and MC3T3 co-culture groups in the 0.85 kPa hydrogels (Figure 2d). Gene expression of OPN was not significantly altered in the culture groups (Figure 2h). ***Tumor-bone cell signaling:*** IL-6 expression was significantly downregulated in the RAW264.7 co-culture and MC3T3 co-culture groups when compared to 4T1 monoculture in the 0.58 kPa hydrogels (Figure 2f). There was no significant change in gene expression of PTHrP between the culture groups for any hydrogel stiffness condition (Figure 2g).

To further understand the effects of these individual biochemical signals on tumor spheroid growth, we extended our computational model to account for these experimentally observed changes in gene expression (TGF-β, TNF-α, and IL-6). Supplementary Table 1 presents the change in the gene expression compared to 4T1 monoculture calculated based on the qRT-PCR gene expression results. We included these values in our computational model and related them to a change in proliferation of the tumor cells (Supplementary Table 2, Supplementary Figure 2). Model calibration (see Section 5.13 and Method 1 in Supplementary Note 1) demonstrated that TNF-α has an inhibitory effect (*γα*= −0.075) on tumor spheroid growth, while IL-6 and TGF-β positively influence growth (*γ*6= 0.125 and *γb*= 0.04, respectively). Overall, our model predicts an inhibitory effect of osteoclast precursors on tumor spheroid growth. In the case of 0.58 kPa hydrogels, we see that this effect stems from an associated increase in TNF-α signaling and a decrease in TGF-β signaling in the presence of osteoclast precursors (Figure 2d). In the absence of osteoclast precursors (i.e. MC3T3+4T1), TNF-α signaling is reduced while TGF-β is increased and tumor spheroids are larger than in the culture groups with RAW264.7 cells (Figure 2b, c, i, j). In the tri-culture group for the same stiffness, these effects were mitigated by high IL-6 signaling and thus larger tumor spheroids were observed in the tri-culture group than in the RAW264.7 co-culture group (Figure 2b, c, i, j). This was also replicated when we took another approach where we capped any change in signal (Δ*S*_*i*_) that was greater than 1, at 1, signifying a saturation of that signal (see Method 2 in Supplementary Note 1, Supplementary Figure 3). We also reported that the spheroids in the MC3T3 co-culture groups (in each stiffness group) experienced higher internal pressure than those in the RAW264.7 co-culture and tri-culture groups (see Supplementary Figure 3b). The predicted compressive spheroid stresses (0.394-0.701 kPa) were largely within the range reported for murine tumors and avascular tumor spheroids (0.4-8.0 kPa)39-41. Taken together, our data suggests that osteoclast precursors have an inhibitory effect on tumor spheroid growth, which is partially mitigated by the presence of osteoblasts.

### 2.3 Development of a Mineralized and Multicellular Bone Model incorporated into a Bioreactor

To more closely mimic the in vivo physiological and mechanical environment of bone tissue, we sought to fabricate a mineralized multicellular model by (1) culturing OCY454 osteocytic cells and MC3T3-E1 cells within gelatin-transglutaminase hydrogels in osteogenic media for 21 days, followed by (2) encapsulating murine RAW264.7 osteoclast precursor cells in gelatin-transglutaminase hydrogel layers onto these mineralized models and (3) applying mechanical compression using a custom built bioreactor (0.5% strain^42^, 1 h per day for 7 days) (Figure 3a).

**Figure 3:**
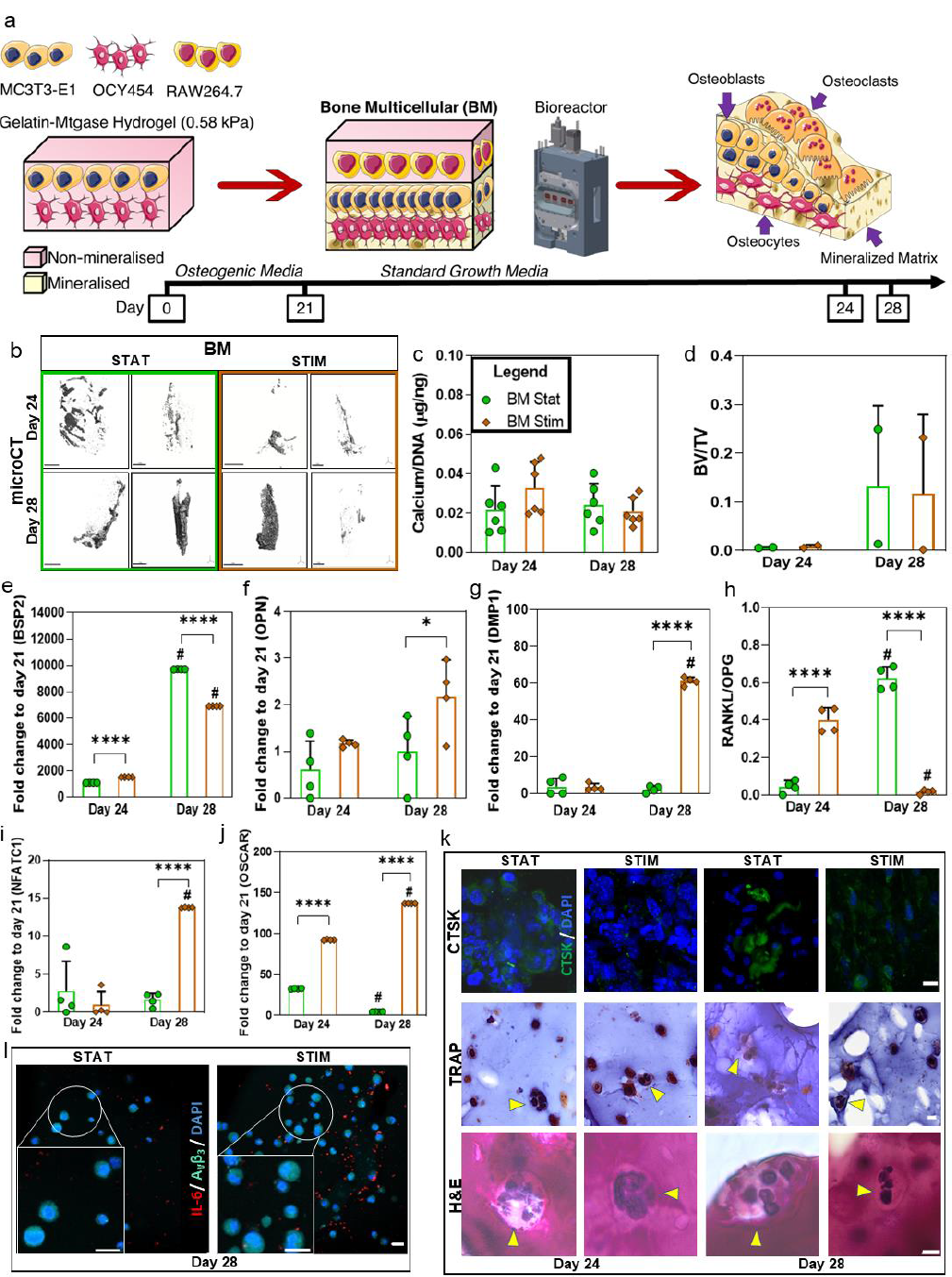
Development of a Bone Multicellular (BM) model with confirmed osteogenic differentiation, mineral deposition, and active osteoclasts. (**a**) OCY454 and MC3T3-E1 cells were encapsulated within gelatin-transglutaminase hydrogels and then cultured in osteogenic media for 3 weeks, at which timea layer of hydrogel encapsulated RAW264.7 cells was added and mechanical compression was applied daily for 7 days. Osteogenic mineralisation was quantified by **(b)** Micro-CT analysis (scale bar: 1 mm), **(c)** a calcium assay and quantitative comparison of the **(d)** bone-like mineral volume fraction (BV/TV).**Osteogenic differentiation** was confirmed by gene expression, relative to day 0: **(e)** BSP2, **(f)** OPN, **(g)** DMP1**. Osteoclast differentiation** was confirmed by gene expression **(h)** RANKL:OPG, **(i)**, NFATC1 **(j)** OSCAR and by **(k)** Immunofluorescence staining for CTSK (scale bar: 10 μm) and histology for TRAP (scale bar: 20 μm) and H&E (scale bar: 10 μm) at day 24 and day 28. **(l)** Immunofluorescence for IL-6 (to assess for inflammation) and α_v_β_3_ (to investigate cellular attachment to the matrix) at day 28 (scale bar: 20 μm, inset scale bar: 20 μm). #: significance compared to day 24, *: p ≤ 0.05, ** : p ≤ 0.01, ***: p ≤ 0.001, ****: p ≤ 0.0001.

We carried out quantitative real-time PCR (qRT-PCR) to understand the changes in osteogenic genes. Under mechanical stimulation, we observed an upregulation of osteogenic genes. Specifically, gene expression for bone sialoprotein (BSP2 – intermediate osteoblast marker) was upregulated when compared to static controls at day 24 (Figure 3e). We also observed an upregulation in Osteopontin (OPN – early osteoblast marker) and dentin matrix acidic phosphoprotein 1 (DMP1 – late osteoblast marker) gene expression under mechanical stimulation when compared to static controls at day 28 (Figure 3f, g), whereas the gene expression of BSP2 was downregulated in mechanically stimulated groups compared to static controls at day 28 (Figure 3e). We also carried out staining for α_v_β_3_, which plays an important role in cellular attachament to the extra-cellular matrix as well as mechanotransduction^43^. Under both stimulated and static conditions we observed positive staining for α_v_β_3,,_ indicative of osteoblast and osteocyte matrix attachments, by day 28 (Figure 3l).

We carried out biochemical analyses to quantify the calcium content (Figure 3c) of our multicellular bone model (BM) and analyzed these constructs for mineral deposition by micro-CT analysis (Figure 3b), as per previous work^44^. A volume of interest (VOI) of 3.95 mm^3^ was segmented using Scanco Medical software (Switzerland) and evaluated for bone-like mineral volume fraction (BV/TV) (Figure 3d). These models demonstrated a high degree of mineral deposition by micro-CT, with a noted increase in mineralization between day 24 and day 28. There was no difference in calcium content under static and stimulated conditions at both day 24 and day 28 (Figure 3c).

Next, we investigated osteoclast activity in these models. We quantified the gene expression ratio of RANKL (receptor activator of NF-κβ ligand) to OPG (Osteoprotegerin), which is an important indicator of the formation of osteoclasts^45, 46^ and observed that this ratio was increased in the mechanically stimulated BM models at day 24, when compared to static controls (Figure 3h). However, we report a significant decrease in the RANKL:OPG ratio under mechanical stimulation at day 28, compared to static controls (Figure 3h). This in keeping with the known inhibitory effect of mechanical loading on bone resorption^47, 48^. We then quantified the expression of Nuclear Factor of Activated T Cells 1 (NFATC1) and osteoclast-associated receptor (OSCAR), which are regulators of osteoclast differentiation^49, 50^, and observed that these two genes were significantly upregulated under mechanical stimulation in the BM model when compared to static controls at day 28 (Figure 3i, j).

To visually confirm the presence of multinucleated active osteoclasts, we then carried out immunofluorescence and histology for Cathepsin-K (CTSK), Tartrate Resistant Acid Phosphatase (TRAP), and Haemo-toxylin and Eosin (H&E). We confirmed the differentiation of osteoclast precursors to multinucleated active osteoclasts (nuclei > 3), which stained positive for CTSK and TRAP, in the multicellular bone model (BM) at days 24 and 28 under both mechanically stimulated and static conditions (Figure 3k).

### 2.4 Metastatic Multicellular Models recapitulates osteolytic behavior

Next, we fabricated metastatic multicellular (MM) models by layering murine RAW264.7 and 4T1 breast cancer, encapsulated in hydrogel layers, onto previously mineralized hydrogels (described above), which were then subjected to mechanical stimulation at 0.5% strain at 1 h per day for 7 days (Figure 4a). We confirmed osteolytic behavior based on increased cancer cell activity, reduced osteoblast activity, increased osteoclastogenesis and pro-osteoclastogenic gene expression. As it is known that tumor cells secrete biochemical factors such as PTHrP and interleukins (e.g. IL-6) during osteolytic metastasis^51^, metastatic behavior in our models was indicated by an increase in the PTHrP gene expression and histological staining for IL-6 in the static MM model at day 28 compared to the BM model (Figure 4i, m). This increased tumorigenic activity stimulated a pro-osteoclastogenic response by means of a significant increase (p ≤ 0.05) in the RANKL: OPG ratio and OSCAR gene expression for the static MM model compared to the static BM model by day 28 (Figure 4j, k).

**Figure 4:**
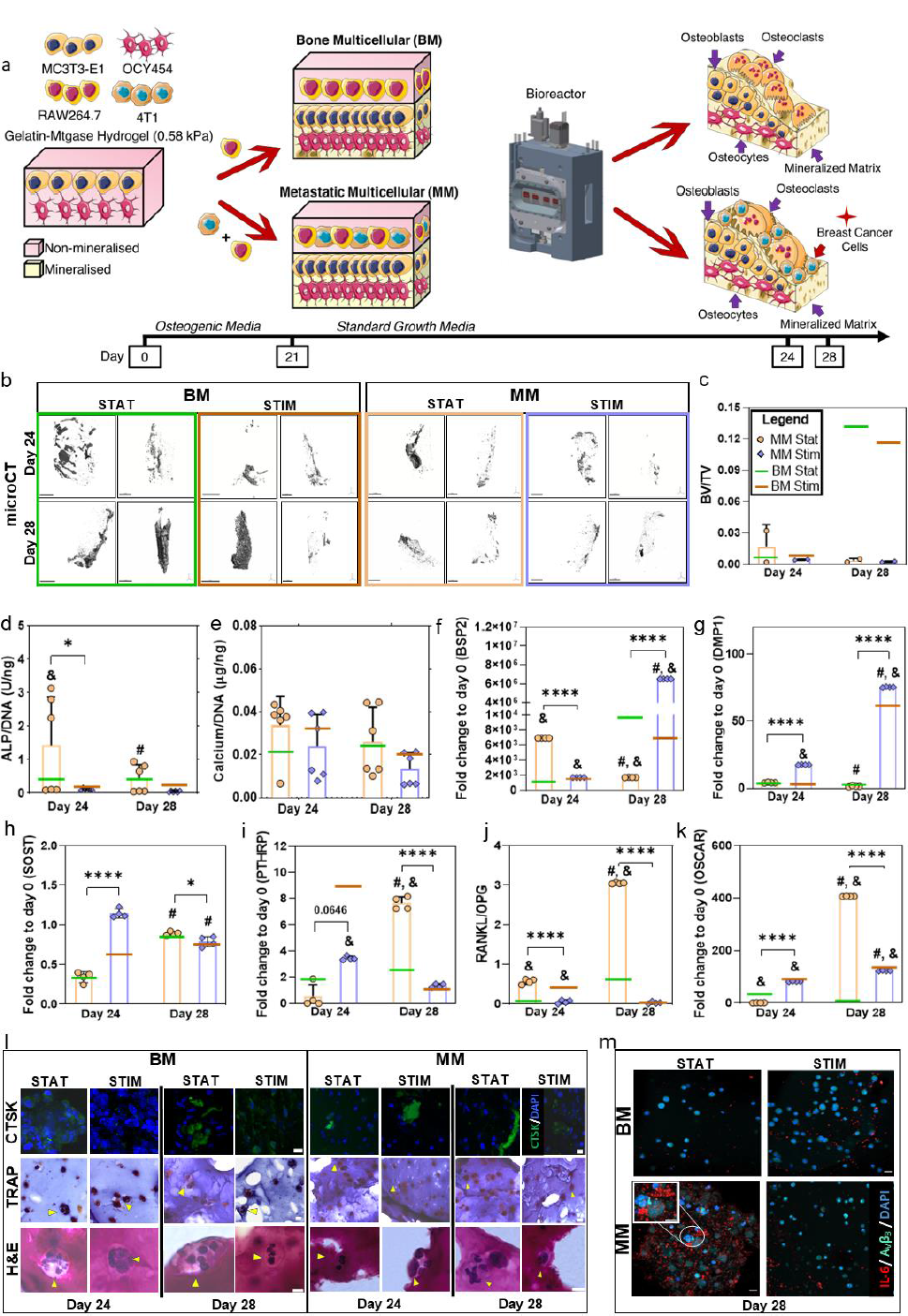
Metastatic Multicellular model exhibits an osteolytic metastatic response, which is attenuated under mechanical compression. (**a**) Schematic of the timeline and the process of encapsulation of OCY454 and MC3T3-E1 cells within gelatin-transglutaminase hydrogels followed by culture in osteogenic media for 3 weeks followed by the addition of either a layer of hydrogel encapsulating RAW264.7 (for BM model) or a layer of RAW264.7 and 4T1 cells (MM Model) in 1:1 ratio and the application of mechanical compression for 7 days. **(b)** Micro-CT analysis of the BM and MM models (scale bar: 1 mm). **(c)** Bone-like mineral volume fraction (BV/TV). **(d)** ALP activity and **(e)** Calcium content in the MM model compared to BM model. Expression of genes, relative to day 0, in the MM model compared to the BM model: **(f)** BSP2, **(g)** DMP1, **(h)** SOST, **(i)** PTHrP, **(j)** RANKL:OPG, **(k)** OSCAR. **(l)** Immunofluorescence staining for CTSK (scale bar: 10 μm) and Histology for TRAP (scale bar: 20 μm) and H&E (scale bar: 10 μm) in the BM and the MM models at day 24 and day 28. **(m)** Immunofluorescence for IL-6 and α_V_β_3_ at day 28 in the BM and the MM models (inset: IL-6 (indicative of inflammation and tumor activity) and α_V_β_3_ (indicative of cellular attachment) positive micrometastasis-like cell aggregates in the MM model under static conditions) (scale bar: 20 μm). &: significance compared to BM model of the corresponding experimental group at the particular timepoint, #: significance compared to day 24, *: p ≤ 0.05, **: p ≤ 0.01, ***: p ≤ 0.001, ****: p ≤ 0.0001.

We investigated how the osteolytic environment influenced osteogenesis (ALP and Calcium) and the expression of osteogenic genes (OPN, BSP2 and DMP1). While ALP activity increased in the presence of breast cancer cells at day 24, when compared to static BM model (p ≤ 0.05, Figure 4d), there was no significant difference between the groups at day 28. No difference in calcium content was observed between the MM and the BM models (Figure 4e). While BSP2 expression was upregulated in the static MM model compared to static BM model at day 24, by day 28 BSP2 expression was downregulated in the static MM model, suggesting reduced osteoblast activity (Figure 4f). Taken together, we observe an osteolytic response in our model due to the presence of breast cancer cells.

### 2.5 Osteolytic metastatic response is attenuated under mechanical compression

We hypothesized that mechanical stimulation would alter the osteolytic response that was captured in our model. Therefore, we characterized metastatic activity in the mechanically stimulated MM models and found that the gene expression of PTHrP and immunostaining for IL-6 were significantly downregulated/decreased when compared to static control at day 28 (Figure 4i, m). We then quantified osteoclast activity in our MM models and report a significant upregulation in the expression of OSCAR for the mechanically stimulated MM model at day 24 compared to static controls. However, we observed a significant downregulation of OSCAR under stimulation at day 28 (Figure 4k) relative to static controls. Most interestingly, mechanical stimulation resulted in a reduction in RANKL:OPG in our MM models at both days 24 and 28, when compared to static controls (Figure 4j). These results together suggest that mechanical compression attenuates the osteolytic metastatic response in our metastatic model.

We finally investigated whether mechanical compression would alter osteogenic activity. We report that ALP activity decreased in the mechanically stimulated MM models compared to the static controls at day 24 (Figure 4d). By day 28, there was no significant difference in ALP activity or calcium content between static and stimulated MM models (Figure 4d, e). While the expression of BSP2 at day 24 was downregulated, we observed an upregulation in BSP2 gene expression at day 28 in the MM models under mechanical stimulation, relative to static controls (Figure 4f). DMP1 and SOST expressions were significantly upregulated in the mechanically stimulated MM model compared to static controls, at both time points for DMP1 and at day 24 for SOST (Figure 4g, h). Expression of SOST at day 28 was downregulated in the mechanically stimulated metastatic groups relative to the static groups (Figure 4h). Together, these results confirm that mechanical compression rescues osteoblast activity from the suppressive effects of the tumor cells.

## 3. Discussion

Bone is subjected to unique physical and biomechanical conditions that are not easily replicated in vitro, but biomaterial-based models are a step closer to achieving more representative models for studying bone pathology. How mechanical cues, arising from growth of metastasized tumor cells within the bone, may elicit mechanobiological responses in both the tumor cells and bone cells, is not clearly understood. Here, we first sought to investigate the influence of matrix stiffness on tumor spheroid growth. Next, we investigated the coupled influence of matrix stiffness and tumor cell-bone cell signaling on tumor spheroid growth. Our results reveal for the first time a synergistic influence of osteoclasts, osteoblasts and 3D matrix stiffness on tumor spheroid growth. Finally, we developed an advanced *in vitro* model that could faithfully mimic the in vivo multicellular, mineralised and mechanical environment. We applied this model for studies of tumor spheroid evolution, tumor cell-bone cell signalling, and osteolytic and pro-osteoclastogenic activity. Our results reveal that mechanical loading can induce breast cancer cell signaling and inhibit osteoclastogenesis and osteoblast activity in early-stage bone metastasis.

There are inherent limitations in using cell lines for our studies, which may behave differently in terms of cell-cell signaling and mechanobiological responses compared to primary human cells. However, isolating sufficient human cells for each cell type and inter-patient variability limited their use, whereas our experiments included mouse bone and mouse cancer cells to avoid potential effects of cross-species interactions. Immune and endothelial cells were not included in our models, which may play an additional role in influencing tumor and bone cell activity during metastasis. However, this study specifically sought to study bone-tumor cell signaling without the complexity of additional cell influences, which could be explored in future studies. We should also note that our study incorporated only compression stimulus, whereas other mechanical stimuli, such as perfusion, may play an important role in stimulating bone and tumor cells in vivo. The mechanical stimulus (0.5% compressive strain) in this study was chosen to induce osteogenic differentiation^42^ and would also be expected to induce perfusion due to the movement of fluid through the nanoscale porosity of the gelatin hydrogel. It is also to be noted that there were only two samples from each group for micro-CT, hence our conclusions regarding osteogenic activity and mineralization are based on biochemical and qRT-PCR analyses. Our models are also not fully representative of in vivo bone, in terms of the extent of mineral deposition and stiffness, which did not approach that of mature bone in vivo (∼0.5-25 GPa). However, it is important to note that the stiffness of our static mineralized hydrogels increased up to 9-fold at day 21 and 24-fold at day 35 compared to day 0, so substantial mineralization of the 3D models in this study was achieved.

Our experimental results and computational model simulations provide a novel understanding regarding the stress-dependent nature of tumor cell proliferation, and how tumour cell-bone cell interactions influence this behaviour. Specifically, through a coupled experimental and computational approach, we show that tumor cell growth within a stiff substrate induces a compressive stress, which reduces tumor cell proliferation (Figure 1g). More importantly, we show for the first time that the co-culture of 4T1 cells with osteoblasts or osteoclast precursors further reduces tumor spheroid size. In particular significant reductions in tumor spheroid size were reported in co-culture with osteoclast precursors and we proposed that this effect is explained by changes in TNF-α and TGF-β1 gene expression (Figure 2j), which encode TNF-α and TGF-β cytokines secreted by RAW264.7 cells that are known to regulate tumor growth^52–57^. Indeed, our computational predictions confirmed the potential role for TNF-α inhibiting tumor spheroid growth. We propose that the lower TNF-α signaling in the presence of osteoblasts alone allowed larger tumor spheroids to form (Figure 2j). Our computational models also predicted that IL-6, an inflammatory cytokine released by both tumor cells and osteoblasts, had a proliferative effect on tumor spheroid growth, which is in keeping with the known effects of IL-6 in terms of promoting tumor growth and suppressing osteoclastogenesis of precursor cells^58–61^, which may, in-turn, affect the production of TGF-β^62^. Although previous mathematical models have been developed and applied to study the interaction between tumor and bone cells, and the signaling pathways involved^63–65^, they have not considered the mechanics of tumor growth. This is the first study that has modelled the combined influence of the tumor cell-bone cell interaction on the growth of tumor spheroids in a confined 3D environment. By incorporating biochemical signaling between bone and the tumor cells and predicted differences in tumor spheroids sizes within hydrogels of the same stiffness, we provided an advanced mechanistic understanding of our experimental results. Specifically, these findings provide evidence that, in addition to compressive stress due to growth, biochemical interactions between the tumor and bone cells modulate tumor spheroid growth.

Our work first confirmed the development of an advanced 3D in vitro bone-like multicellular model, which was well mineralized and expressed known osteogenic markers. Previous in vitro models aiming to mimic a multicellular bone-like microenvironment have utilized multiple cell types, including human umbilical endothelial cells, bone marrow mesenchymal stem cells, osteoblasts and osteoclasts encapsulated within fibrin based hydrogels^34, 66^. More recently, 3D in vitro models have captured paracrine signalling between human primary cells, osteocytes embedded in collagen gels, and osteoclasts^67^ and osteoblasts^68^ grown on a porous membrane, in co-culture ^69^ and have recreated osteoblast-osteoclast remodeling of mineralized silk fibroin based bone-mimicking template^70^. However, in vivo the bone microenvironment is highly mineralized and experiences continuous mechanical loading arising from daily physical activity and osteocytes mediate bone cell resorption and formation accordingly. Hence it is essential to include mechanical loading and osteocytes within a mineralized niche to account for the in vivo microenvironment and the influence of mechanobiological processes on bone cell activity. Additionally, in these previous studies, osteoclastic differentiation of mononuclear cells or monocytes has been achieved through the use of exogenous cytokines (RANKL or M-CSF). It is important to note that our model was capable of inducing differentiation of osteoclast precursors towards multinucleated active osteoclasts without the need for any externally added cytokines. In this way, we demonstrate the potential for these primed mineralized models to be used for further studies of osteoclast precursor differentiation and activation. In contrast to a previous study in our laboratory, which demonstrated that the application of mechanical stimulation resulted in increased mineralization in osteoblast-only encapsulated hydrogels in vitro^44^, calcium content was similar for the bone-like models cultured with and without mechanical stimulation. Mechanical stimulation of osteoblasts activates specific signaling pathways^71^, which stimulate the production of anabolic growth factors and synthesis of extracellular matrix proteins and mineral^72^. In fact, dynamic compression can promote the differentiation and mineralization of both osteoblasts and stem cells toward osteogenesis both in vitro and in vivo^73–77^. It may be that here the inclusion of other cells in the multicellular niche suppressed the typical induction of mineral production by osteoblasts cultured in isolation under mechanical stimulation. In particular, the inclusion of osteoclasts may have inhibited mineral production by signaling to osteoblasts. Of relevance, semaphorin 4D produced by osteoclasts inhibits bone formation^78^ in a manner similar to that of OPG, which also limits the osteoclast-promoting actions of RANKL. In this study, OPG was significantly upregulated in the mechanically stimulated bone-like model compared to the static controls. Another explanation may be that the active osteoclasts degraded the mineral during the timeframe of the study. Indeed, pro-osteoclastogenic gene expression (increased RANKL:OPG ratio) by osteoblasts and osteocytes was evident and the expression of osteoclast differentiation markers (NFATC1, OSCAR) were significantly greater in the mechanically stimulated bone-like groups compared to static.

Next, we implemented this approach towards the successful development of a bone metastasis model. Previous studies have developed advanced 3D in vitro models of bone metastasis by cellularizing bio-printed scaffolds with human bone cells, with a mineralized microenvironment, and then incorporating prostate or breast cancer cells^32, 33^,or by encapsulating breast-cancer cells (MCF-7, MDA-MB-231) in a hydrogel and studying interactions with human primary osteoblasts in a separate cryogel^54^. A recent study developed a bone metastasis model by incorporating human bone cells (osteoblasts, osteoclasts and macrophages) with endothelial cells and MDA-231 breast cancer cells within fibrin hydrogels, which was applied to investigate the efficacy of drug treatments (doxorubicin,rapamycin) to attenuate the growth of cancer cells^34^. Osteocyte paracrine signaling has been shown to inhibit metastatic breast cancer cell growth, whereas fluid flow-stimulated osteocytes reverse this effect and cause increased tumor cell migration and proliferation^79, 80^. Fluid shear stress-stimulated osteocytes cause a reduction in tumor cells migration and lead to an increase in apoptosis via signaling to osteoclasts^81^. Thus, the mechanical environment is also a key regulator of bone and tumour cell activity, whichhas not been incorporated into previous models of bone metastasis. With this in mind, this study addresses this gap in the development of advanced 3D in vitro models by including a multicellular niche within a mineralized environment and by including mechanical signals through a custom bioreactor.

Our multicellular model of bone metastasis recapitulated the increased secretion of PTHrP and IL-6 by tumor cells, which we propose explains the increase in expression of RANKL (by osteoblasts and osteocytes) and induced osteoclast differentiation and activity (Figure 5a). In fact, IL-6 enhances osteoclastogenesis by upregulating RANKL in osteocytic MLO-Y4 cells^82^. Overall, upregulated expression of these metastatic, pro-osteoclastogenic, and osteoclast markers in our metastatic model are in keeping with the known osteolytic nature of 4T1 breast cancer cells and confirms the ability of our experimental model to recapitulate the osteolytic metastatic process^83^. The initial increase in ALP activity and gene expression of BSP2, a marker of osteoblast mineralization^84^, in the bone metastasis model under static conditions may be attributed to 4T1 cells, which can express these markers as an early response to the presence of a mineralized matrix^85^. Interestingly, tumor cells can assume a phenotype similar to bone cells^86^ in order to survive and proliferate better in bone tissue. Alternatively, the presence of the cancer cells might induce an osteogenic response by osteoblasts. Indeed, disseminated tumor cells have been shown to predominantly colonize the osteogenic niche in vivo, with active canonical WNT signaling and newly formed osteoblasts in the vincinty of the osteolytic lesions^87^.

**Figure 5:**
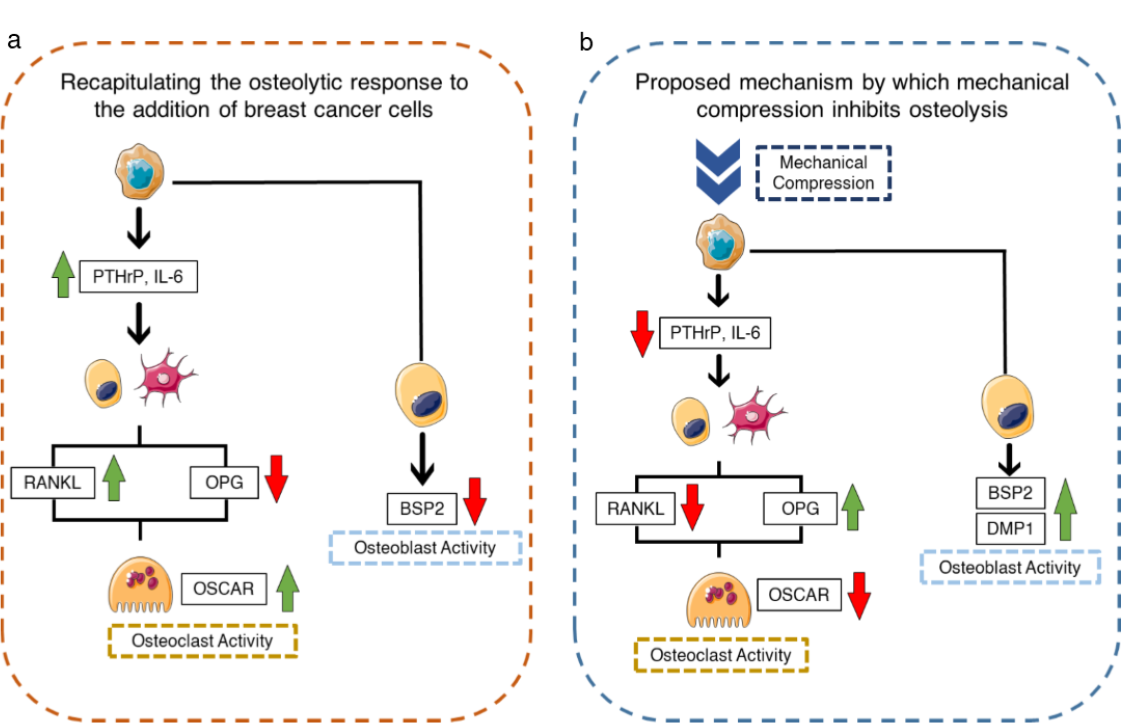
**(a) Recapitulating the osteolytic response to the addition of breast cancer cell using the 3D in vitro multicellular mineralized model.** Breast cancer cells secrete biochemical factors that stimulate osteoclastogenesis through the RANKL/OPG pathway and downregulate osteoblastic activity. **(b) Proposed mechanism by which mechanical compression inhibits this osteolytic response.** Mechanical compression leads to a reduction in the expression of osteolytic factors, which in turn leads to a reduction in expression of the factors supporting osteoclastogenesis, while genes associated with osteoblast activity are upregulated.

On the basis of our findings, we propose that mechanical compression can reduce the expression of metastatic genes, reverse pro-osteoclastogenic activity and rescue suppressed osteogenic activity (Figure 5b). Previous studies have demonstrated that mechanical stimulation can prevent breast cancer associated-osteolysis by driving osteogenic activity^38^, reducing tumor cell proliferation^37^ and reducing osteocyte hypoxia and apoptosis^88^. The anti-catabolic effects of mechanical loading may also be modulated by nitric oxide (NO) and mechanical stimulation stimulates the release of NO from bone cells^89, 90^, which induces a decrease in the RANKL-OPG ratio. In our model, we report an early upregulation in sclerostin expression in the metastatic model under mechanical stimulation, which was accompanied by a downregulation of BSP2 and upregulation of OSCAR, which provide a possible mechanism for the inhibitory effects of mechanical stimulation on osteoclastogenesis in our model. Interestingly, sclerostin released by metastatic breast cancer cells reduces osteoblast differentiation in vitro, whereas sclerostin inhibition reduced bone destruction in an animal model of breast cancer-bone metastasis^91^. Thus, our model provides an advanced understanding of the interactions between tumor and bone cells, and the role of the mechanical environment for regulating the vicious cycle. These models will be used in the future to study how inhibition of mechanobiological responses can attenuate tumor cell-bone cell signaling and the development of osteolytic bone metastases.

## 4. Conclusions

This study provides an advanced understanding of how tumor growth and osteolysis are regulated by the evolving mechanical environment during tumor invasion of bone. Using novel experimental and computational tumor growth models we showed, for the first time, a synergistic influence of osteoclasts, osteoblasts and 3D matrix stiffness on tumor spheroid growth. By developing an advanced multicellular and mineralized model, we successfully captured the osteolytic-metastatic process and revealed that mechanical stimulus could provide protective effects against tumor-induced osteolysis. Our model of early-stage bone metastasis may be utilized to test the efficacy of mechanobiological inhibitors for preventing bone and tumor cell activation associated with bone metastasis.

## 5. Materials and Methods

### 5.1 Expansion of cells

OCY454 osteocytic cells (Centre for Skeletal Research Bone Cell Core, an NIH-funded program (P30AR075042), which is supported by NIAMS) were expanded in cell culture flasks coated with type I collagen (0.15 mg/ml in 0.02 M acetic acid). The growth media used was α-MEM, which was supplemented with 10% fetal bovine serum (FBS), 100 U/mL penicillin, 100 μg/mL streptomycin and 2 mM L-glutamine. The cells were then incubated at a permissive temperature of 33°C, in an incubator at 5% CO_2_. For differentiating OCY454 cells, a semi-permissive temperature of 37°C was chosen. MC3T3-E1 osteoblastic cells (ATCC, USA) were expanded in standard growth media (as for OCY454 cells). RAW264.7 osteoclast precursors (ATCC, USA) were expanded in low glucose DMEM supplemented with 10% FBS (HyClone), 100 U/mL penicillin, 100 μg/mL streptomycin and 2 mM L-glutamine. Murine mammary carcinoma 4T1 cells were expanded in media (Roswell Park Memorial Institute - RPMI) supplemented with 10% FBS, 100 U/mL penicillin and 100 μg/mL streptomycin. MC3T3-E1, RAW264.7 and 4T1 cells were expanded at 37°C, in an incubator with 5% CO_2_. Cell culture media was replenished every 3 days.

### 5.2 Three-dimensional (3D) Static Culture - Spheroid Growth

Cell encapsulating gelatin hydrogels of three different stiffnesses (0.58 kPa, 0.85 kPa,1.1 kPa) were developed by crosslinking equal volumes of 9% cell laden gelatin and nano-Hydroxyapatite (nHa) (5.5% v/v) with sterile microbial transglutaminase (mtgase) containing Activa WM (Aginomoto Foods) of different concentrations (0.3%, 0.6%, 1% w/w). These hydrogels were used to encapsulate separate layers of 4T1, MC3T3-E1, and RAW264.7 cells in various culture configurations (Monoculture: 4T1, Co-culture: RAW+4T1, MC3T3+4T1, Tri-culture: MC3T3+RAW+4T1). For the Tri-culture group MC3T3 cells were suspended in 9% gelatin and enzymatically cross-linked with different concentrations of mtgase (0.3%, 0.6% and 1.0% w/w of gelatin) and nHA (1:1:1 ratio), to yield a final gelatin concentration of 3% w/v, a nHA concentration of 5.5% v/v and an MC3T3 cell density of 2×10^6^ cells/mL. These hydrogel suspensions were pipetted into custom-made polydimethylsiloxane (PDMS) wells in a layer (1 mm (height) x 4 mm (breadth) x 13 mm (length)) and cooled at 4°C for 8 minutes to encapsulate MC3T3 cells with separate individual hydrogel layers of varying stiffness (0.58 kPa, 0.85 kPa, 1.1 kPa). Similarly, RAW264.7 cells were suspended in 9% gelatin, cross-linked with the different concentrations of mtgase and nHA (1:1:1 ratio) and pipetted on top of the hydrogel layer of MC3T3 cells of the corresponding stiffness and allowed to gel at 4°C for 8 minutes. A PKH26 Red Fluorescent (λ_ex_= 551 nm; λ_em_= 567 nm) Cell Linker Kit (Sigma-Aldrich, Ireland) was used to label the cell membrane of the 4T1 cells. These labelled 4T1 cells were then suspended in gelatin and cross-linked with different concentrations of mtgase with nHa to yield hydrogel suspensions of the three different stiffness (as previously described) and then pipetted on top of the previously deposited hydrogel layer of RAW264.7 cells of the corresponding stiffness and cooled to gel at 4°C for 8 minutes.

RAW264.7 co-cultures (RAW + 4T1) and MC3T3 co-cultures (MC3T1+4T1) were created by suspending the first cell type in gelatin, crosslinking with different concentrations of mtgase with nHa, pipetting into PDMS wells and cooling at 4°C for 8 minutes before repeating the process for the second cell type in the co-culture and layering on top of the previously gelled layer of cells of the corresponding compression modulus (Figure 2a). The cell number was kept at 100,000 cells/hydrogel layer (2×10^6^ cells/mL of hydrogel). The media for the co- and tri-culture experiments was formulated by maintaining an equal ratio (1:1 or 1:1:1) of MC3T3, RAW 264.7, and 4T1 growth media. The media was changed every 2 days for the duration of the culture period. All hydrogel groups were cultured for 7 days (Fig. 1a, Fig. 2a). The tumor spheroid sizes (section 5.9), DNA content (section 5.6.1) and expressions of genes implicated in tumor cell-bone cell signaling (PTHrP, IL-6, TGF-β TNF-α and OPN) were quantified (section 5.12).

### 5.3 Fabrication of Bilayered models for mineralization

OCY454 cells were suspended in gelatin with an initial concentration of 9%. This cell suspension was cross-linked with mtgase and with nHA (1:1:1 ratio), to yield a final gelatin concentration of 3% w/v, mtgase concentration of 0.3% w/w of gelatin, a nHA concentration of 5.5% v/v, and an OCY454 cell density of 2×10^6^ cells/mL. To fabricate the OCY454 layer, this hydrogel suspension was then pipetted into the PDMS wells (1.33 mm (height) × 6 mm (breadth) × 17 mm (length)) and allowed to cool at 4°C for 8 min. Next, an MC3T3 gelatin/nHA suspension was cross-linked with mtgase, as described for OCY454 cells. This suspension was pipetted on top of the OCY454 layer and cooled at 4°C for 8 min to form an osteoblast layer (1.33 mm (height) × 6 mm (breadth) × 17 mm (length)). These bilayered models were cultured in standard OCY 454 growth media (α-MEM, 10% FBS, 100 U/mL penicillin, 100 μg/mL streptomycin, 2 mM L-glutamine) supplemented with osteogenic factors (50 μM ascorbic acid, 10 mM β-glycerophosphate, and 100 nM dexamethasone). These models were cultured for 21 days to enable mineralization. Media was replenished every 3 days.

### 5.4 Fabrication of bone multicellular (BM) and metastatic multicellular (MM) models

To fabricate the bone multicellular (BM) model, RAW264.7 encapsulated gelatin/nHA suspension was cross-linked with mtgase as described previously. This suspension was pipetted on top of the osteoblast layer of the bilayered models, which had been cultured for 21 days as described in 5.3 above. These were allowed to cool at 4°C for 8 min.

To fabricate the metastatic multicellular (MM) model, RAW264.7 and 4T1 (1:1) encapsulated gelatin/nHA was enzymatically cross-linked with mtgase as described previously. This suspension was pipetted on top of the osteoblast layer of the bilayered models and allowed to cool at 4°C for 8 min. These models were cultured in the following growth media formulations: (1) Bone Multicellular (BM): RAW 264.7 growth media and MC3T3/OCY454 growth media mixed in a ratio of 1:2 and (2) Metastatic Multicellular (MM): RAW 264.7 growth media, 4T1 growth media and MC3T3/OCY454 growth media mixed in a ratio of 1:1:2. Culture media was replenished every 3 days for the culture period to allow for proliferation and differentiation of the pre-osteoclasts and breast cancer cells (7 days).

### 5.5 Mechanical Stimulation

Multicellular models (BM and MM) from each experimental group were transferred from well plates to a sterile custom-built compression bioreactor (VizStim) containing their respective cell culture media, as described previously. Four samples from each group were placed within each bioreactor chamber. These constructs were subjected to compression at 0.5 % strain for 1 h per day for 7 days. At day 3 and day 7, the constructs were removed from culture and cut into quarters in both the sagittal plane and the transverse planes. These quarters were then processed for biochemical (see section 5.6), histological (see section 5.7), immunofluorescence (see section 5.10), micro-computed tomography (see section 5.11) and qRT-PCR (5.12) analyses. Static control groups were also included for all experimental conditions.

### 5.6 Biochemical Analyses

#### 5.6.1 DNA Assay

Constructs were rinsed twice using phosphate buffered saline (PBS), weighed and then stored at −80°C. For the assay, the constructs were thawed and digested in 3.88 U/mL papain in 0.1 M sodium acetate, 5 mM L-cysteine-HCl, 0.05 M EDTA, at pH of 6.0 and a temperature of 60°C under rotation for 18 h. DNA content was measured using the Hoechst 33258 DNA assay, as described previously ^92^.

#### 5.6.2 Extracellular ALP activity

Alkaline phosphatase (ALP) secreted into the culture media was measured for each experimental group. At each time point, constructs were kept in the incubator for 2 h after the last bout of mechanical stimulation. The media was then removed and stored at −80°C. ALP activity was measured through a colorimetric assay of enzyme activity (SIGMAFAST p-NPP Kit), where p-nitrophenyl phosphate (pNPP) (nmol) acts as a phosphatase substrate and the ALP enzyme as a standard. After thawing, 40 μL of the culture medium was transferred to a 96-well plate in triplicate with 50 μL of pNPP solution. The well plate kept away from direct light at room temperature for 1 h. Colorimetric readings were taken at 405 nm using a microplate reader (Synergy HT Multi-mode). Using a standard ALP curve, the measured readings were converted to ALP activity. ALP production was normalized to DNA content to confirm whether variation in mineralization potential were dependent on the overall mineralization capacity of individual cells.

#### 5.6.3 Mineralization

Calcium content in the mineralized multicellular constructs was evaluated using a Calcium Liquicolour kit (Stanbio Laboratories, Syntec, Ireland) as per the manufacturer’s protocol. Constructs were rinsed with PBS, weighed and stored at −80°C. For analysis, constructs were thawed and digested in 1 mL of 1M hydrochloric acid (HCl) and at 60°C under rotation for 18 h. For each of the digested models and assay standards 10 μL was added to a 96-well plate in triplicate followed by 200 μL of the working solution. Absorbance readings were obtained at 550 nm using a microplate reader, as previously described^93^.

### 5.7 Histology

Constructs from each group were fixed in 4% paraformaldehyde (PFA) overnight, equilibrated in 30% sucrose solution, embedded in OCT embedding medium (VWR), sectioned at 20μm using a Cryostat (CM1850, Leica) and affixed to microscope slides (Fisher Scientific). The sections were stored at −20°C. Sections were then allowed to thaw and rinsed with PBS. Sections were stained with Haematoxylin and Eosin (H&E) and TRAP (all Sigma Aldrich). For H&E, sections were immersed in Mayer’s Hematoxylin, followed by immersion in running tap water. Slides were immersed three times in HCl/ethanol solution (1% HCl in 70% ethanol) before being stained with Eosin (Thermo scientific) for 3 minutes. Slides were then washed again with several deionized water rinses. Of note, only cells with three nuclei or more were considered to be osteoclasts. For TRAP, sections were rinsed with deionized water and stained for TRAP activity using a commercial kit and counterstained with Mayer’s Hematoxylin for 10 seconds before being immersed in running tap water. All sections were air-dried before mounting with DPX. Images were acquired using a light microscope (BX 43 Olympus Microscope, software cellSens).

### 5.8 Cell Viability

Cell viability within 4T1 monoculture constructs was assessed by carrying out LIVE/DEAD staining. Constructs were transferred to a 24-well plate at day 5 of culture and washed with PBS before staining. A staining solution containing 500 µl of PBS, 8 µM calcein AM (Biocambridge Sciences, BT80011-1) and ethidium homodimer-1 (EthD-1) (Biocambridge Sciences, BT40014) was added to the samples for 2 hours at 37 °C in the dark. All images were captured using an Olympus IX50 inverted fluorescence microscope.

### 5.9 Fluorescent Staining and Imaging

Fluorescent staining was performed on hydrogels from 3D static culture spheroid growth experimental groups to visualize the actin cytoskeleton and nuclei of the cells, and the formation of 4T1 spheroids. Hydrogels were fixed on days 3 and 7 using 4% paraformaldehyde at 4°C overnight. Cells within these hydrogels were then permeabilized with 0.5% Triton-X in PBS for 10 mins at 4°C under agitation. Samples were stained with phalloidin-FITC at 1.25 μg/mL (1:400) to stain the actin cytoskeleton and DAPI dilactate (1:2000) to identify cell nuclei, respectively. Z-stack imaging was carried out using a Fluoview FV1000 confocal laser scanning microscope system (Olympus) at a magnification of 20x (oil) with a step size of 5 μm. All stacks were obtained at the same intensity setting between groups. NIH ImageJ software was used to analyze stacks of images at maximum intensity projections.

### 5.10 Immunofluorescent Staining

Mineralized multicellular BM and MM Constructs were stained for DAPI, Cathepsin K (CTSK), Interleukin-6 (IL-6), α_V_β_3_, PTHrP, dentin matrix acidic phosphoprotein 1 (DMP1) and OPN. At day 24 and 28 of culture, constructs were fixed in 4% paraformaldehyde at 4°C overnight. The cells within the constructs were then permeabilized using 0.5% Triton-X in PBS for 10 min at 4°C under agitation. Actin cytoskeleton and nucleus were stained using phalloidin-FITC at 1.25 μg/mL (1:400) and DAPI dilactate (1:2000), respectively.

For CTSK staining, the constructs were blocked in 1% bovine serum albumin (BSA) for 1h under agitation. Then these samples were incubated in Alexa Fluor 488 conjugated mouse monoclonal anti-CTSK antibody (1:100) at 4°C overnight (Merck) and then for 1h under agitation at room temperature. Constructs were further counterstained with DAPI dilactate (1:2000) to visualize the nucleus. For IL-6, samples were incubated with goat polyclonal IL-6 antibody (Santa Cruz) (1:100) at 4°C overnight. After rinsing in 1% BSA, the constructs were incubated with Donkey anti goat Alexa 647 (Jackson ImmunoResearch) for 1 h under agitation at room temperature. For, integrin α_v_β_3_ staining, anti-α_v_β_3_ antibody directly conjugated to Alexa Fluor® 488 (1:100) (Santa Cruz) was utilized. Constructs were further counterstained with DAPI dilactate (1:2000) and rinsed again with 1% BSA solution.

Maximum intensity projections were obtained from z-stacks taken at 20x (oil lens) magnification on a Fluoview FV1000 confocal laser scanning microscope (Olympus) with a step size of 5 μm between each slice. NIH ImageJ software was used to measure the overall fluorescence intensity of the stained images. Fluorescent intensities were calculated by subtracting the background integrated density from respective integrated density.

### 5.11 Micro-Computed Tomography (μCT)

Constructs were fixed in 4% paraformaldehyde at 4°C overnight for micro-computed tomography (μCT). Constructs from both static and mechanically loaded groups were evaluated for mineralization by μCT scanning (Scanco Medical μCT100) using the following profile: a 9 mm holder, 45 kVp tube voltage, 200 μA intensity, Al filter, 5 μm voxel size, 300 ms integration time, and a frame averaging value of 2. A density threshold of 370 mg HA/cm^3^ (3964 Hounsfield units) was applied to exclude hydrogel and yet retain the mineralized regions, as per previous work ^44^. A volume of interest (VOI) of 3.95 mm^3^ was segmented using Scanco Medical software (Switzerland) and evaluated for mineralization properties such as bone-like mineral volume fraction (BV/TV).

### 5.12 Quantitative Real-Time PCR

Total RNA content was extracted using a custom Cetyl trimethylammonium bromide (CTAB) method ^94^. Each construct was digested in 250 μL of RNAse-free proteinase K solution (5 mg/mL) containing 5 mM CaCl_2_. The digested constructs were mixed with 500 μL of CTAB solution (2% [w/v], 1.4 M NaCl, 20 mM EDTA, and 100 mM Tris, pH 8) with 1% (v/v) β-mercaptoethanol. Then 500 μL of chloroform was added to each construct and centrifuged at 14,000 g for 2 min at room temperature. The upper/aqueous phase was transferred to a fresh tube and 800 μL of isopropanol (Fisher) was added to precipitate total RNA by centrifugation (14,000 g, 15 min, room temperature). The pellet was washed with 600 μL ethanol (70% v/v) and dissolved in 20 μL RNAse-free water for 15 min at 65°C. RNA purity and yield were assessed using a spectrophotometer (DS-11 FX, DeNovix), with 260/280 and 260/230 ratios over 1.8 for all samples. Then, 500 ng of RNA from each sample of static culture spheroid growth model and 300 ng of RNA from each sample of mineralized multicellular model were transcribed into cDNA using Qiagen Quantinova reverse transcription kits and thermal cycler (5PRIMEG/O2, Prime). Relative gene expression was studied by quantitative real-time polymerase chain reaction (qRT-PCR). The genes of interest included osteoblast and osteocyte specific genes: bone sialoprotein (BSP2) and osteopontin (OPN), dentin matrix acidic phosphoprotein 1 (DMP1) and sclerostin (SOST), osteoclast specific genes: nuclear factor of activated T cell 1 (NFATC1) and osteoclast associated receptor (OSCAR), genes associated with bone resorption: Receptor Activator for Nuclear Factor κ B Ligand (RANKL) and osteoprotegerin (OPG), and genes implicated in metastasis: Parathyroid hormone-related protein (PTHrP), Interleukin 6 (IL-6), Transforming growth factor β (TGF-β) and Tumor necrosis factor alpha (TNF-α). Rplp0 and Rpl13A were used as reference genes (Table 1).

**Table 1.**
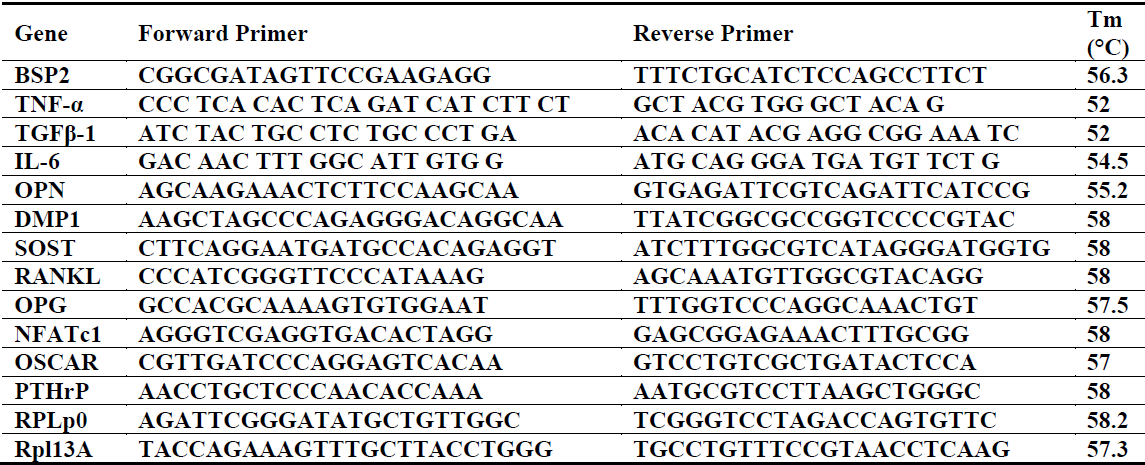
List of primers employed for qRT-PCR, including sequences (5’-3’) and annealing temperatures (Tm).

qRT-PCR was carried out using a Qiagen Quantinova SYBR Green PCR kit and a Quant Studio 5 PCR machine (Applied Biosciences). The PCR was conducted with an enzyme activation step at 95°C for 2 min, and then for 40 cycles with the following steps: 5 s denaturation at 95°C and specific annealing temperatures for each primer pair (Table 1) for 18 s. Analysis of the results was done using the Pfaffl method (Pfaffl, 2001), and the results are expressed as relative quantitative changes to 4T1 monoculture samples of the corresponding hydrogel stiffness at day 3 of culture for the 3D static spheroid growth experimental groups. For the multicellular mineralized BM and MM models, results are expressed as relative quantitative changes to samples from day 0.

### 5.13 Computational Model Development

In developing a model to predict the growth of a tumor spheroid, we first considered that the number of tumor cells (*c*_*t*_) is governed by the rate of cell proliferation (*k*_*p*_) and apoptosis (*k*_*a*_), such that

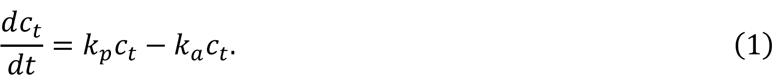

It has been proposed that the size of tumor spheroids is sensitive to compressive stresses imposed by matrix expansion ^36, 95^, likely a compound effect of suppression of cell proliferation and cell compaction. Mechanical inhibition of cell division may be considered through a dependency in the proliferation rate *k*_*p*_, such that

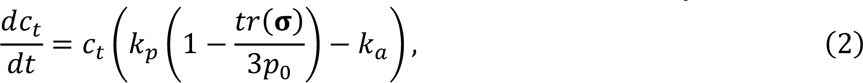

where ***σ*** is the Cauchy stress tensor at a given point in a growing spheroid, and *p*_0_ is a reference pressure for mitotic inhibition. Although the division rate is also dependent on access to nutrients and solutes (i.e. oxygen, glucose), in this study, the growth of small (∼60 *μm*) spheroids is being analysed and therefore it is assumed that they are uniformly perfused. Equation (2) may be further simplified such that

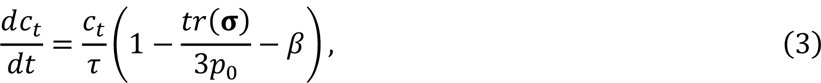

where *τ* = 1/*k*_*p*_ and *β* = *k*_*a*_/*k*_*p*_. *β* (= 0.077) was obtained by calculating 4T1 cell proliferation and death rates based on data provided in literature ^96, 97^. This equation follows a form similar to the model proposed by Ambrosi et al (2017) ^70^. To analyse the co-culture and tri-culture of breast cancer cells with osteoclast precursors and osteoblasts, we further extended this model by considering how gene expressions of TNF-α, TGF-β, and IL-6 (*S*_*α*_, *S*_*β*_, and *S*_6_, respectively) can promote or inhibit tumor spheroid growth. The model accounted for a change in expression of these genes relative to the baseline 4T1 monoculture, 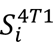, such that Δ*S*_*i*_ = *S*_*i*_ − 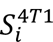 (see Supplementary Table 1) from gene expression levels are directly taken from the experimental qRT-PCR data measured in this study (Figure 2d-f). Upregulated and downregulated signals from the gene expression experimental data were accounted for based on the following criteria: Δ*S*_*i*_ < −0.5 (strongly down-regulated), Δ*S*_*ri*_ = −1; −0.5 ≤ Δ*S*_*i*_ ≤ −0.25 (weakly downregulated), Δ*S*_*ri*_ = −0.5; −0.25 < Δ*S*_*i*_ < 0.25 (negligible change), Δ*S*_*ri*_ = 0; 0.25 ≤ Δ*S*_*i*_ ≤ 0.5 (weakly upregulated), Δ*S*_*ri*_ = 0.5; 0.5 < Δ*S*_*i*_ (strongly upregulated), Δ*S*_*ri*_ = −1. The associated values are reported in Table 2. Following from this, the rate of tumor cell proliferation may be given as:

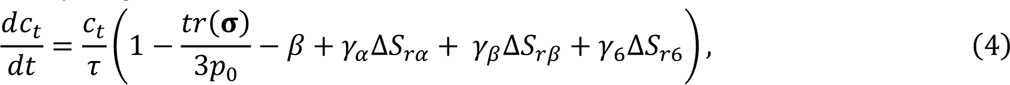

**Table 2.**
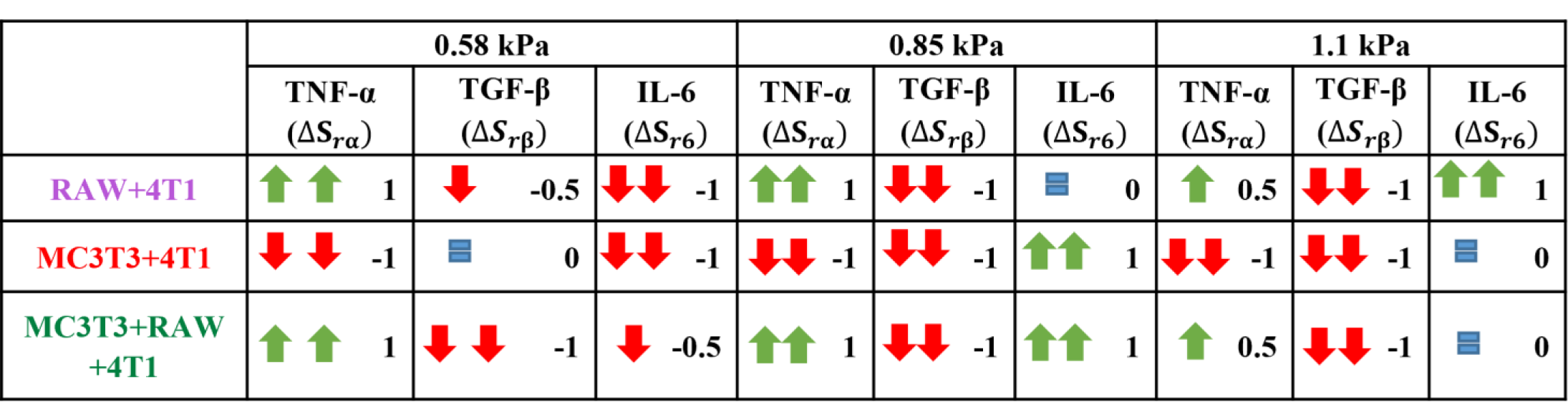
Signal values: Δ*S*_*rα*_ (TNF-α), Δ*S*_*rβ*_ (TGF-β), and Δ*S*_*r*6_ (IL-6) for each culture condition.

where *γ*_*α*_, *γ*_*β*_ and *γ*_6_ are factors that relate the changes in signaling to the change in proliferation. Clearly, a positive value of *γ*_*i*_ suggests that the associated biochemical factor promotes tumor cell proliferation while a negative value reduces proliferation.

#### 5.13.1 Finite Element Modelling of Tumor Spheroid Growth

A 2D axisymmetric model of a single tumor cell, with a radius of 7.5 µm, was generated using finite element (FE) software Abaqus (2019), whereby the cell was embedded within a hydrogel of dimensions 300 µm x 150 µm. The distance between the cell boundary and the outer surface of the hydogel (142.5 µm) was chosen to minimise edge effects. The geometry was meshed using 4-noded axisymmetric elements (CAX4) with a total of 1368 elements. Boundary conditions were applied to the distal edges of the hydrogel to prevent axial and radial displacement, and at the line of symmetry to prevent displacement in the x-axis direction.

The tumor cell proliferation model was implemented in Finite Element (FE) software Abaqus (2019) via a user-defined material subroutine (UMAT). To simulate growth, a multiplicative decomposition of the deformation gradient **F** into **F**_*g*_ (growth tensor) and **F**_*e*_ (elastic tensor)^95, 98^ was adopted, such that **F** = **F**_*e*_**F**_*g*_. Here, **F**_*g*_ = *λ*_*g*_**I**, where *λ*_*g*_ = (*c*_*t*_/*c*_0_)^1/3^ is the growth stretch dependent on the number of tumor cells *c*_*t*_, **I** is the second-order identity tensor and *c*_0_ is the initial number of cells in a spheroid. From this, the elastic component of the deformation gradient **F** could be obtained from 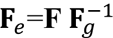. The passive elastic behaviour of the spheroid and hydrogel is described by a Neo-Hookean formulation, with a Cauchy stress given by:

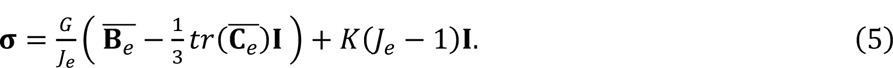

Here, *J*_*e*_ is the determinant of the elastic component of the deformation gradient, 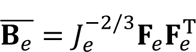 and 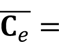 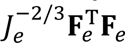 are the left and the right Cauchy-green tensors, respectively. *G* is the material shear modulus, and *K* is the material bulk modulus. Final parameter values obtained from calibration based on experimental results are shown in Supplementary Table 2. Results from parameter sensitivity analysis are displayed in the Supplementary Figure 5.

### 5.14 Statistical Analysis

Statistical analyses were performed using GraphPad Prism software (version 8.4.3). One-way ANOVA with Tukey’s post-hoc test was used to compare the significance in spheroid size and qRT-PCR for the co-culture and the tri-culture experiments. Two-way ANOVA was used for analysis of variance with Bonferroni’s post-hoc tests to compare between different timepoints and experimental groups in all the other experiments. The results are displayed as mean ± standard deviation, with significance at a level of p ≤ 0.05. Three to four replicates were analyzed for each experimental group. The experiments were performed three times.

## Acknowledgments

This research was funded by Irish Research Council Laureate Award Programme 2017/18, MEMETic, IRCLA/2017/217. The authors acknowledge the facilities and scientific and technical assistance of the Centre for Microscopy and Imaging at the National University of Ireland Galway (www.imaging.nuigalway.ie). The authors would also like to acknowledge Servier Medical Art (www.servier.com) for their image bank used to produce the figures.

## Author Contributions

Conceptualization, S.M.N., V.K., E.M., L.McN.; methodology, S.M.N., V.K., A.S.K.V., E.M., L.McN.; validation S.M.N., V.K.; formal analysis, S.M.N., V.K. A.S.K.V.; investigation, S.M.N. V.K.; data curation, S.M.N. V.K.; writing—original draft preparation, S.M.N. V.K., E.M., L.McN; writing—review and editing, S.M.N., V.K., A.S.K.V., E.M., L.McN; visualization, S.M.N., V.K.; supervision, E.M., L.McN; project administration and funding acquisition, L.McN. All authors have read and agreed to the published version of the manuscript.

