## Supplementary File for "A novel mechanobiological model of bone metastasis reveals that mechanical stimulation inhibits the pro-osteoclastogenic effects of breast cancer cells"

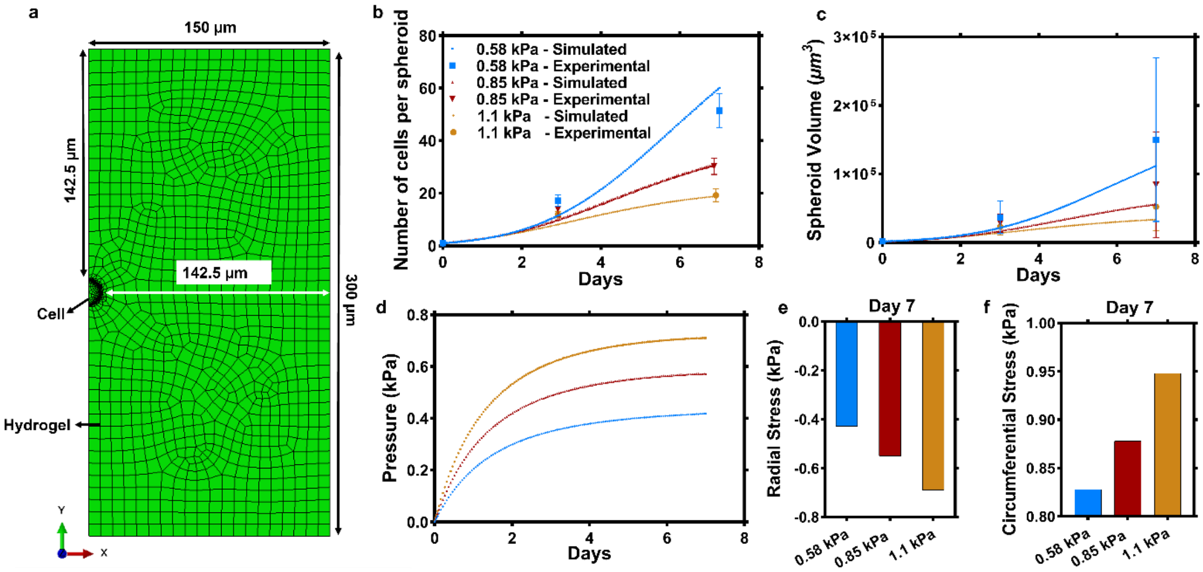


**Supplementary Figure 1:** Mechanical loading reduces tumour spheroid size by restricting cell proliferation. **(a)** Finite element model of a tumor cell embedded within a hydrogel, where the distance between the cell boundary and the outer surface of the hydrogel (142.5 µm) was chosen to minimise edge effects**.** Computational predictions of **(b)** the number of nuclei per spheroid and **(c)** spheroid volume over time area mapped against the experimental results. **(d)** Pressure developed uniformly within these growing spheroids as a response to compressive stress imposed by the substrate. **(e, f)** Radial and circumferential stress developed by day 7 within the hydrogels due to spheroid growth.

| | 0.58 kPa | | | 0.85 kPa | | | 1.1 kPa | | |
|---|---|---|---|---|---|---|---|---|---|
| | TNF-α ($\Delta S_\alpha$) | TGF-β ($\Delta S_\beta$) | IL-6 ($\Delta S_6$) | TNF-α ($\Delta S_\alpha$) | TGF-β ($\Delta S_\beta$) | IL-6 ($\Delta S_6$) | TNF-α ($\Delta S_\alpha$) | TGF-β ($\Delta S_\beta$) | IL-6 ($\Delta S_6$) |
| RAW+4T1 | 1.60 | -0.27 | -0.71 | 0.78 | -0.57 | 0.22 | 0.28 | -0.80 | 0.65 |
| MC3T3+4T1 | -0.85 | 0.12 | -0.65 | -0.62 | -0.60 | 1.12 | -0.66 | -0.67 | -0.16 |
| MC3T3+RAW +4T1 | 3.84 | -0.61 | -0.46 | 2.78 | -0.71 | 0.68 | 0.50 | -0.52 | -0.18 |

**Supplementary Table 1:** Parameters ($\Delta S_i$) depicting change in gene expression compared to 4T1 monoculture, which were calculated based on the qRT-PCR gene expression results (from Figure 2 d-f): $\Delta S_\alpha$ (TNF-α), $\Delta S_\beta$ (TGF-β), and $\Delta S_6$ (IL-6).

### Supplementary Note 1. Coupled biochemical signalling and tumor spheroid growth model calibration

In both our in vitro and in silico models, compressive stresses generated by hydrogel deformation increase with increasing hydrogel stiffness (in the absence of any biochemical interaction). This will lead to higher spheroid pressure in each of the co-culture and tri-culture groups, which in turn will result in a reduction in proliferation (per equation 3). However, individual biochemical signals from the osteoclast precursors and osteoblasts will also alter the growth of the tumor cells. As described in the manuscript (Method 1), we incorporate this signaling in our model through a $\Delta S_{ri}$ value and the associated regulation potential $\gamma_i$, as shown in Supplementary Figure 2. A positive $\Delta S_{ri}$ value denotes an increase in the signal and negative value depicts a decrease in the signal, relative to 4T1 monoculture).

**Method 1:**From the qRT-PCR data obtained in this study, we calculated the $\Delta S_i = S_i - S_i^{4T1}$ values for TNF-α, TGF-β, and IL-6 ($\Delta S_\alpha$, $\Delta S_\beta$, and $\Delta S_6$) corresponding to each culture group (Supplementary Figure 2a, Supplementary Table 1). Depending on the degree of upregulation or a downregulation from the experimental gene expression data (compared to the monoculture), these changes were then accounted for by assuming $\Delta S_i$ as either 0 (negligible change), ±0.5 (slight change) or ±1 (large change) (positive sign in case of upregulation and a negative sign in the case of downregulation) (Supplementary Figure 2b). These values $\Delta S_{r\alpha}$ (TNF-α), $\Delta S_{r\beta}$ (TGF-β), and $\Delta S_{r6}$ (IL-6) were then input into equation 4 (Supplementary Figure 2c). We simulated each of the 9 experimental groups (three hydrogel stiffness and three culture group per stiffness) until good agreement was achieved with the in vitro results (Supplementary Figure 2d). For the simulations, in addition to the calibrated values of: $G_H$, $K_H$, $G_C$, $K_C$, $\tau$, $P_o$, and $\beta$, the single set of calibrated values for $\gamma_\alpha$, $\gamma_\beta$ and $\gamma_6$ are shown in Supplementary Table 2. A parameter sensitivity analysis was performed with results shown in Supplementary Figure 5. We were also able to replicate these trends when we took another approach (Method 2 described below, Supplementary Figure 3)

| Parameters | Values |
|---|---|
| Shear moduli of hydrogels ( $G_H$) | 0.22 kPa, 0.32 kPa, and 0.42 kPa, respectively |
| Bulk moduli of hydrogels ($K_H$) | 0.48 kPa, 0.71 kPa, and 0.92 kPa, respectively |
| Shear Modulus of the Spheroid ($G_C$) | 0.340 kPa |
| Bulk Modulus of the Spheroid ($K_C$) | 23.53 kPa |
| Timescale for tumor cell proliferation ($\tau$) | 0.89 days |

| | |
|---|---|
| Reference pressure for mitotic inhibition ($p_0$) | 2.20 kPa |
| The ratio of apoptosis rate to proliferation rate ($\beta$) | 0.077 [1,2] |
| TNF-$\alpha$ signaling constant ($\gamma_\alpha$) | -0.075 |
| TGF-β signaling constant ($\gamma_\beta$ ) | 0.04 |
| IL-6 signaling constant ($\gamma_6$) | 0.125 |

**Supplementary Table 2:** Parameters used in **Method 1** for calibrating and simulating the computational tumor spheroid growth model.

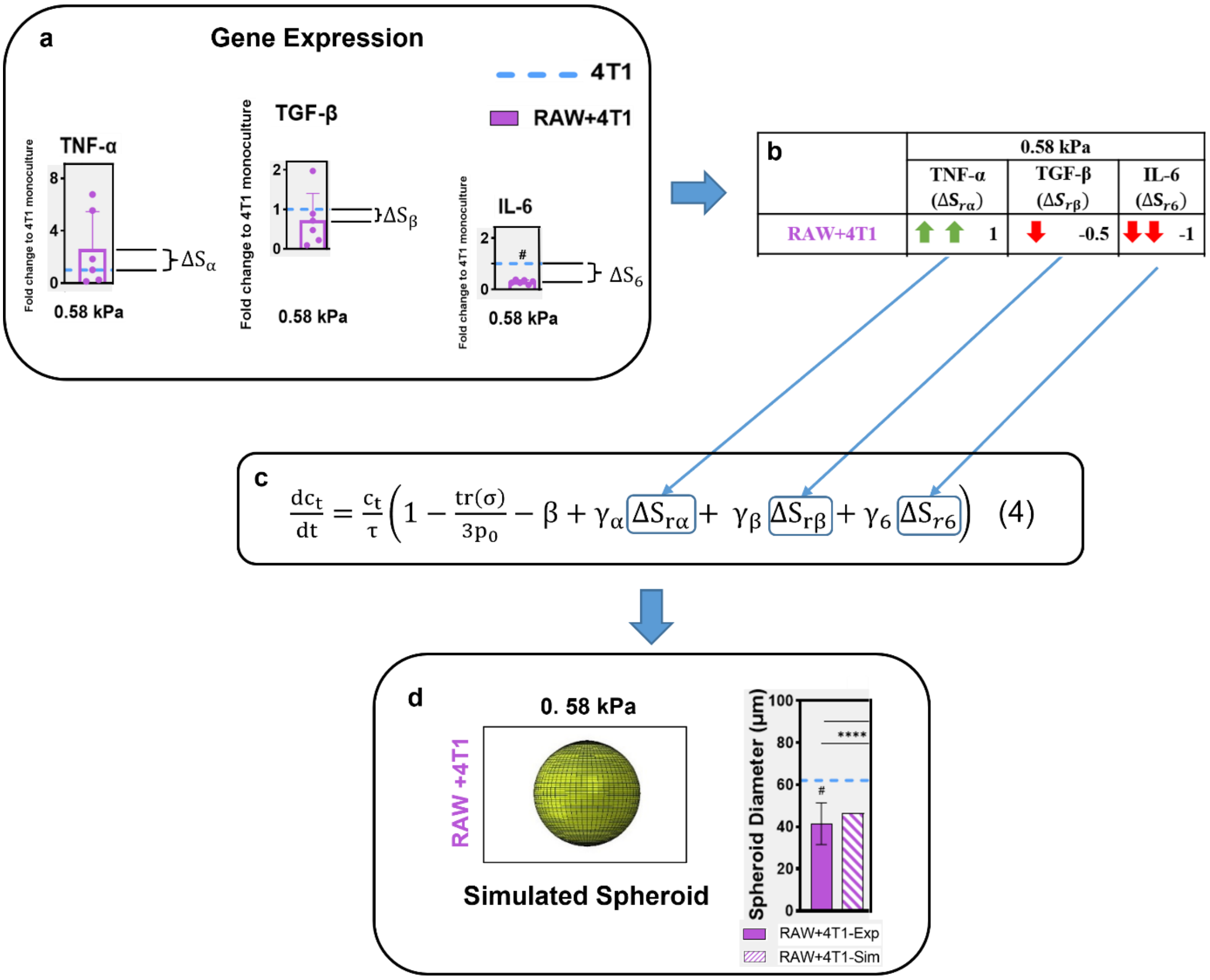


$$\frac{dc_t}{dt} = \frac{c_t}{\tau}\left(1 - \frac{tr(\sigma)}{3p_0} - \beta + \gamma_\alpha \Delta S_{r\alpha} + \gamma_\beta \Delta S_{r\beta} + \gamma_6 \Delta S_{r6}\right) \quad (4)$$

**Supplementary Figure 2: (a)** Schematic describing the process of determining input signals from our experimental gene expression data: the values for $\Delta S_\alpha$ (TNF-α), $\Delta S_\beta$ (TGF-β), and $\Delta S_6$ (IL-6) were determined based on the results of qRT-PCR analysis for a specific culture group and stiffness condition (e.g. RAW + 4T1 and 0.58 kPa). **(b)** These values are then converted to 0, ±0.5 or ±1 based amount on a criteria described in section 5.13 of the manuscript and **(c)** sent as input into equation 4 of our computational model, along with the parameters described in Supplementary Table 1, for a particular set of $\gamma_\alpha$, $\gamma_\beta$, and $\gamma_6$. The model simulations were then conducted for each experimental group per one set of $\gamma_\alpha$, $\gamma_\beta$, and $\gamma_6$ until a good agreement with the in vitro results was achieved. **(d)** An example simulation for tumor spheroid growth for a specific culture group and stiffness condition (RAW + 4T1 and 0.58 kPa) after calibration. This calibration was achieved across all the culture groups and hydrogels.

**Method 2:** From the qRT-PCR data obtained in this study, we calculated the $\Delta S_i = S_i - S_i^{4T1}$ values for TNF-α, TGF-β, and IL-6 ($\Delta S_\alpha$, $\Delta S_\beta$, and $\Delta S_6$) corresponding to each culture group (Supplementary Figure 2a, Supplementary Table 1). Any calculated $\Delta S_i$ value > 1, was capped at 1, signifying a saturation of that signal. (Supplementary Table 3). The resulting new signal values $\Delta S_{r\alpha}$ (TNF-α), $\Delta S_{r\beta}$ (TGF-β), and $\Delta S_{r6}$ (IL-6) were then input into equation 4. We simulated each of the 9 experimental groups (three hydrogel stiffness and three culture group per stiffness) until a good agreement was achieved with the in vitro results (Supplementary Figure 3b). For the simulations, in addition to the calibrated values of: $G_H$, $K_H$, $G_C$, $K_C$, $\tau$, $P_o$, and $\beta$, the single set of calibrated values for $\gamma_\alpha$, $\gamma_\beta$ and $\gamma_6$ are shown in Supplementary Table 4.

| | **0.58 kPa** | | | **0.85 kPa** | | | **1.1 kPa** | | |
|---|---|---|---|---|---|---|---|---|---|
| | **TNF-α ($\Delta S_\alpha$)** | **TGF-β ($\Delta S_\beta$)** | **IL-6 ($\Delta S_6$)** | **TNF-α ($\Delta S_\alpha$)** | **TGF-β ($\Delta S_\beta$)** | **IL-6 ($\Delta S_6$)** | **TNF-α ($\Delta S_\alpha$)** | **TGF-β ($\Delta S_\beta$)** | **IL-6 ($\Delta S_6$)** |
| **RAW+4T1** | **<u>1</u>** | **-0.27** | **-0.71** | **0.78** | **-0.57** | **0.22** | **0.28** | **-0.80** | **0.65** |
| **MC3T3+4T1** | **-0.85** | **0.12** | **-0.65** | **-0.62** | **-0.60** | **<u>1</u>** | **-0.66** | **-0.67** | **-0.16** |
| **MC3T3+RAW +4T1** | **<u>1</u>** | **-0.61** | **-0.46** | **<u>1</u>** | **-0.71** | **0.68** | **0.50** | **-0.52** | **-0.18** |

**Supplementary Table 3:** Signal values: $\Delta S_{r\alpha}$ (TNF-α), $\Delta S_{r\beta}$ (TGF-β), and $\Delta S_{r6}$ (IL-6) for each culture condition in **Method 2**. Signals capped at 1 are underlined.

| **Parameters** | **Values** |
|---|---|
| Shear moduli of hydrogels ( $G_H$) | 0.22 kPa, 0.32 kPa, and 0.42 kPa, respectively |
| Bulk moduli of hydrogels ($K_H$) | 0.48 kPa, 0.71 kPa, and 0.92 kPa, respectively |
| Shear Modulus of the Spheroid ($G_C$) | 0.340 kPa |
| Bulk Modulus of the Spheroid ($K_C$) | 23.53 kPa |
| Timescale for tumor cell proliferation ($\tau$) | 0.89 days |
| Reference pressure for mitotic inhibition ($p_0$) | 2.20 kPa |
| The ratio of apoptosis rate to proliferation rate ($\beta$) | 0.077 [1,2] |
| TNF-$\alpha$ signaling constant ($\gamma_\alpha$) | -0.05 |
| TGF-β signaling constant ($\gamma_\beta$ ) | 0.02 |
| IL-6 signaling constant ($\gamma_6$) | 0.15 |

**Supplementary Table 4:** Parameters used in the **Method 2** for calibrating and simulating the computational tumor spheroid growth model.

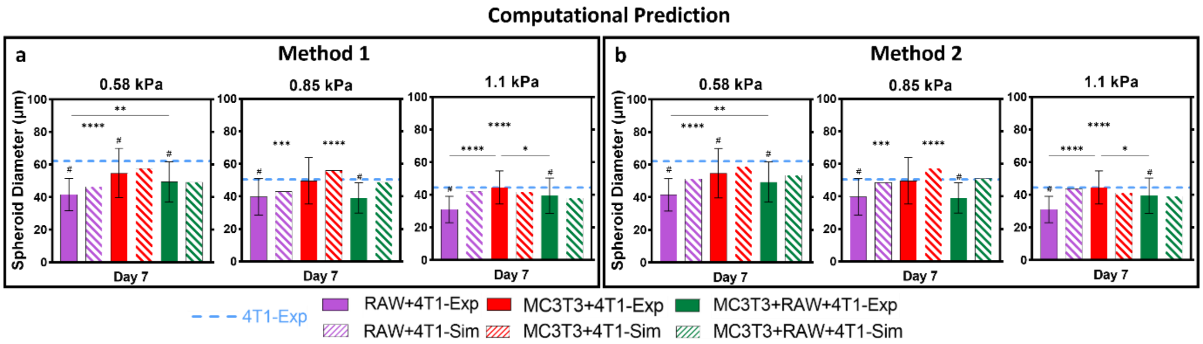


**Supplementary Figure 3:** Comparison of spheroid diameters, plotted against the experimental values (Exp), obtained from the tumor spheroid growth model calibration (Sim) using **(a) Method 1** and **(b) Method 2.**

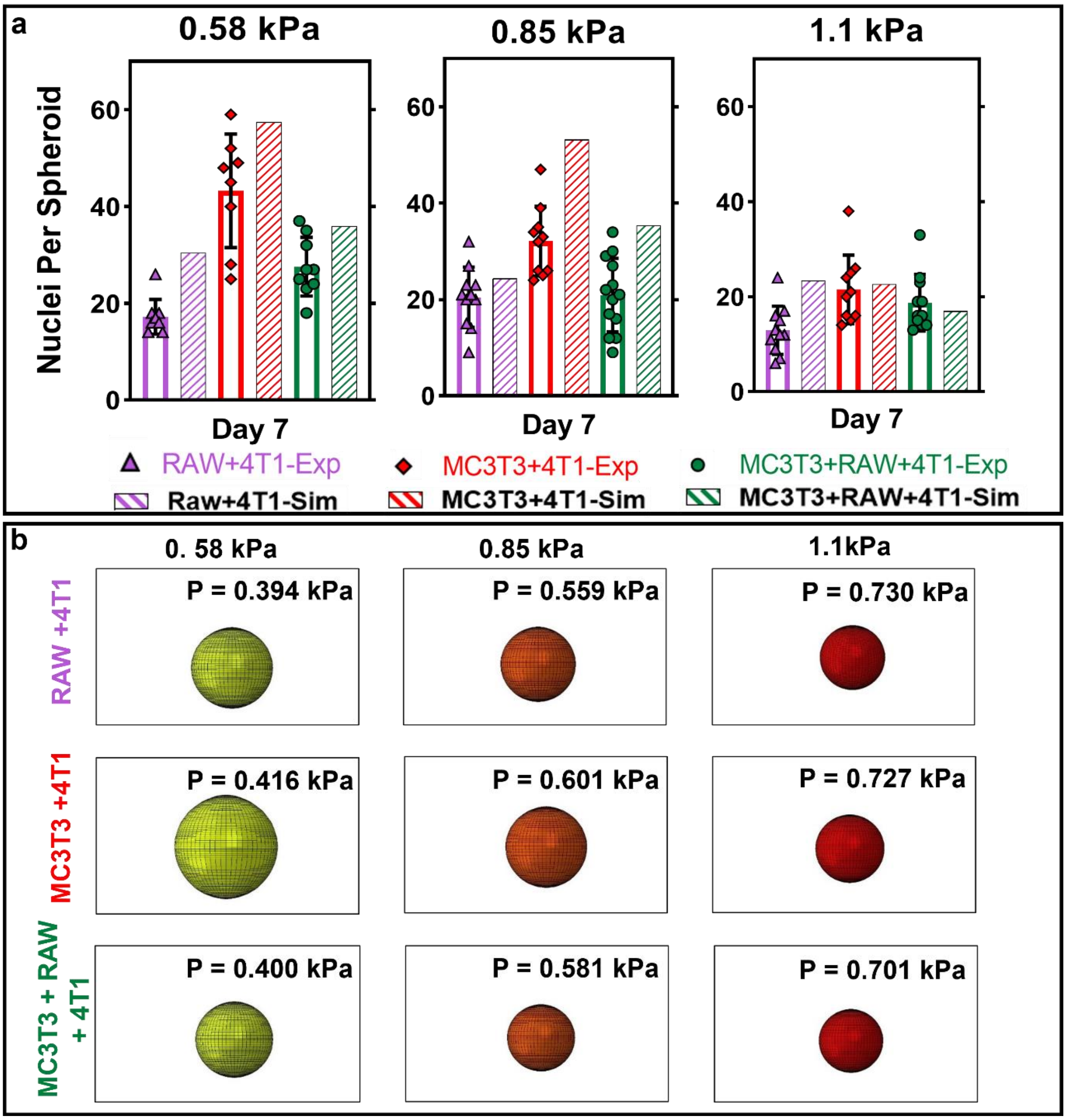


**Supplementary Figure 4: Osteoclast precursors have an inhibitory effect on tumor spheroid growth which is partially alleviated in the presence of osteoblasts (Method 1): (a)** Comparison of nuclei per spheroid at day 7 in the in vitro model with results of simulation from computational tumor growth model (Sim) within the hydrogels of stiffness 0.58 kPa, 0.85 kPa and, 1.1 kPa. **(b)** Simulations predicted that pressure (P) developed uniformly within the growing spheroids as a response to signaling from the bone cells and compressive stress imposed by the hydrogel.

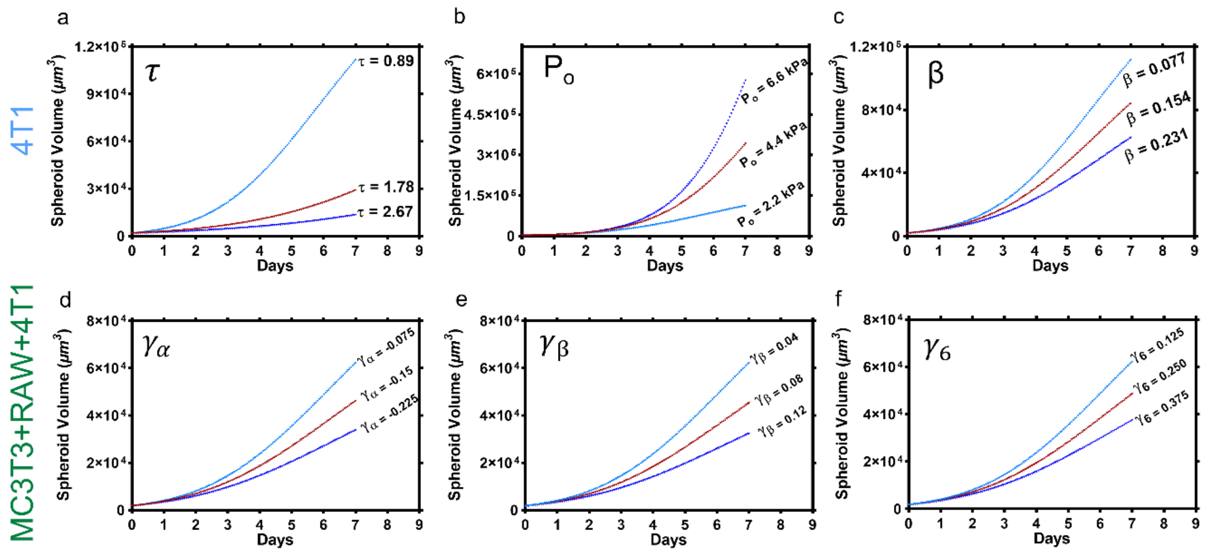


**Supplementary Figure 5: Sensitivity analysis for parameters used in the computational tumor growth model (Method 1): (a)** timescale for tumor cell proliferation ($\tau$), **(b)** reference pressure for mitotic inhibition ( $p_0$), **(c)** ratio of apoptosis rate to proliferation rate ($\beta$), factors relating change in signaling to the change in proliferation for: **(d)** $\gamma_\alpha$ (TNF-α), **(e)** $\gamma_\beta$ (TGF-β) and **(f)** $\gamma_6$ (IL-6) (0.58 kPa hydrogel case displayed here for the 4T1 mono-culture and the tri-culture groups).

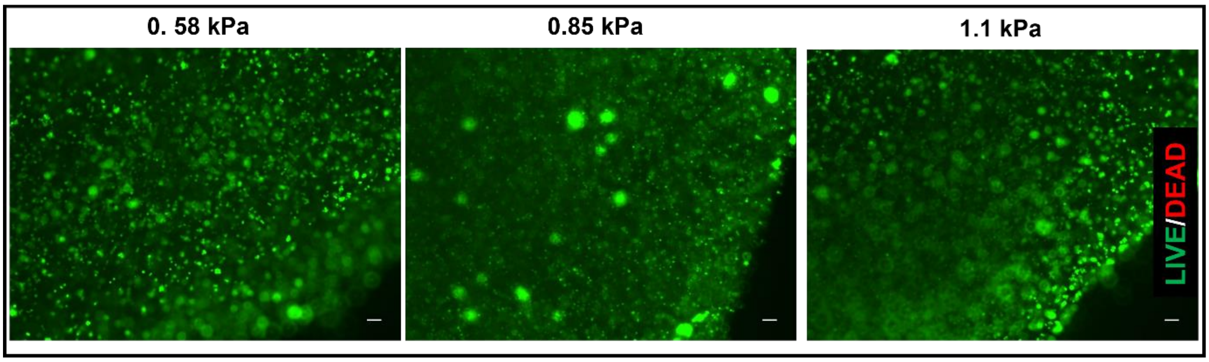


**Supplementary Figure 6:** Representative Live (Green)/Dead (Red) image of 4T1 cells in monoculture encapsulated within hydrogels of different compression moduli (0.58 kPa, 0.85 kPa and 1.1 kPa), at day 5 of culture (Scale Bars: 60 µm).

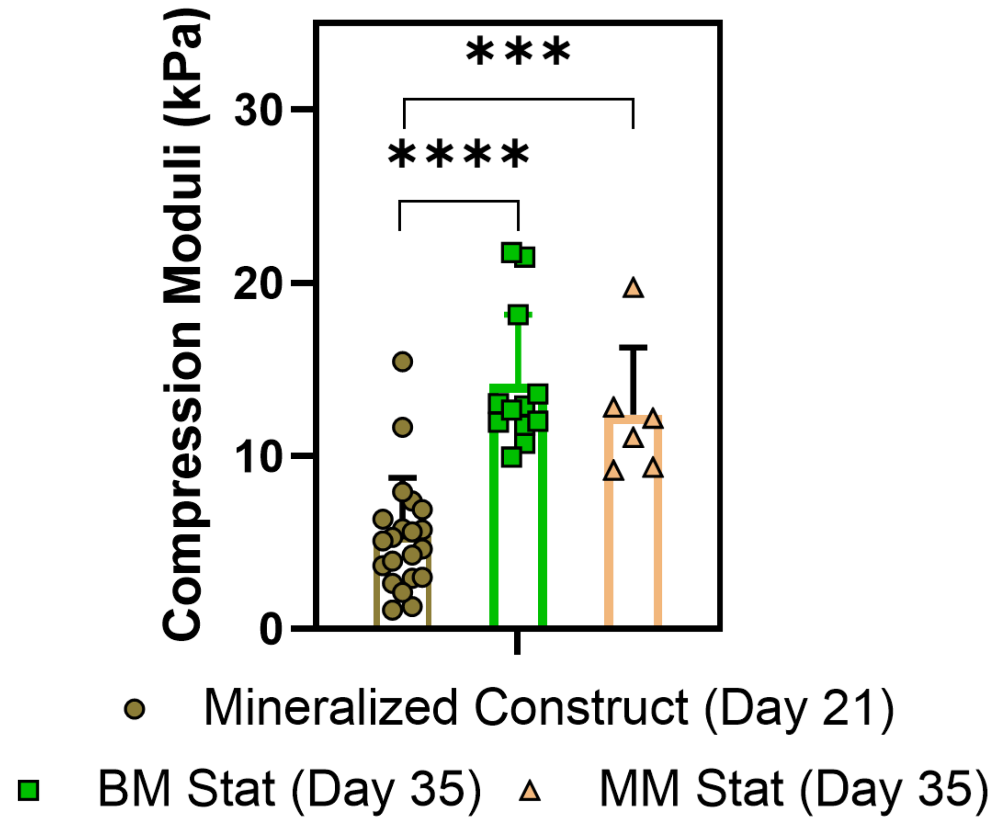


**Supplementary Figure 7: Mineralization in the 3D models results in compression moduli increasing up to 24-fold:** compression modulus of the hydrogels at the end of osteogenic supplementation (day 21) and of the BM and the MM models under static conditions (day 35).
